# Mechanistic machine learning enables interpretable and generalizable prediction of prime editing outcomes

**DOI:** 10.64898/2026.02.20.706353

**Authors:** Alvin Hsu, Peter J. Chen, Angus H. Li, Colin F. Hemez, Xin D. Gao, Markus Terrey, Charlie Nelson, Vijay Selvam, Ana Cristian, Amber N. McElroy, Benjamin J. Steinbeck, Gandhar K. Mahadeshwar, Smriti Pandey, Zachary Barsdale, Paul Z. Chen, Alexander A. Sousa, Holt A. Sakai, Rachel A. Silverstein, Ilias Morad, Ryan K. Krueger, Max W. Shen, Benjamin P. Kleinstiver, Cathleen M. Lutz, Jakub Tolar, Bruce R. Blazar, Mark J. Osborn, David R. Liu

## Abstract

Although prime editing (PE) can effect virtually any specified local change to genomic DNA in living systems, its efficient application currently requires extensive optimization of prime editing guide RNA (pegRNA) sequences. We present OptiPrime, a machine learning model of PE efficiency based on our current understanding of the mechanism of prime editing. OptiPrime achieves state-of-the-art accuracy on PE efficiency prediction and also enables prediction of nicking guide RNA (PE3) and dual pegRNA (twinPE) outcomes. We validated that OptiPrime has learned the determinants of mammalian mismatch repair (MMR), and is therefore well suited for nominating MMR-evasive silent edits that improve PE efficiency. We demonstrate the utility of OptiPrime in a variety of prospective therapeutic contexts, including in primary human and mouse cells. Finally, we show how OptiPrime can be used to achieve highly streamlined and efficient *in vivo* correction of a pathogenic mutation in the brain of a mouse model of *KIF1A*-associated neurological disorder.

## Introduction

Precision gene-editing technologies have revolutionized research in the life sciences, as well as the treatment of both heritable and non-heritable diseases in the clinic^1–48^. Prime editing (PE) is a versatile gene-editing method that uses a programmable nickase and a prime editing guide RNA (pegRNA) to nick a target DNA sequence at a specified location, reverse transcribe the edited sequence from the pegRNA onto the nicked target DNA strand, and guide the cell through DNA repair processes to replace the original DNA sequence and make the resulting edit permanent on both DNA strands^49^ (Figure 1A). PE can install virtually any substitution, small insertion, small deletion, or combination thereof in the genome of a living cell without requiring double-stranded breaks (DSBs) or a donor DNA template (Figure 1B). For example, PE has been used clinically to correct a 2-bp deletion in *NCF1*, resulting in the effective treatment of chronic granulomatous disease (CGD) in multiple patients^50^.

**Figure 1.**
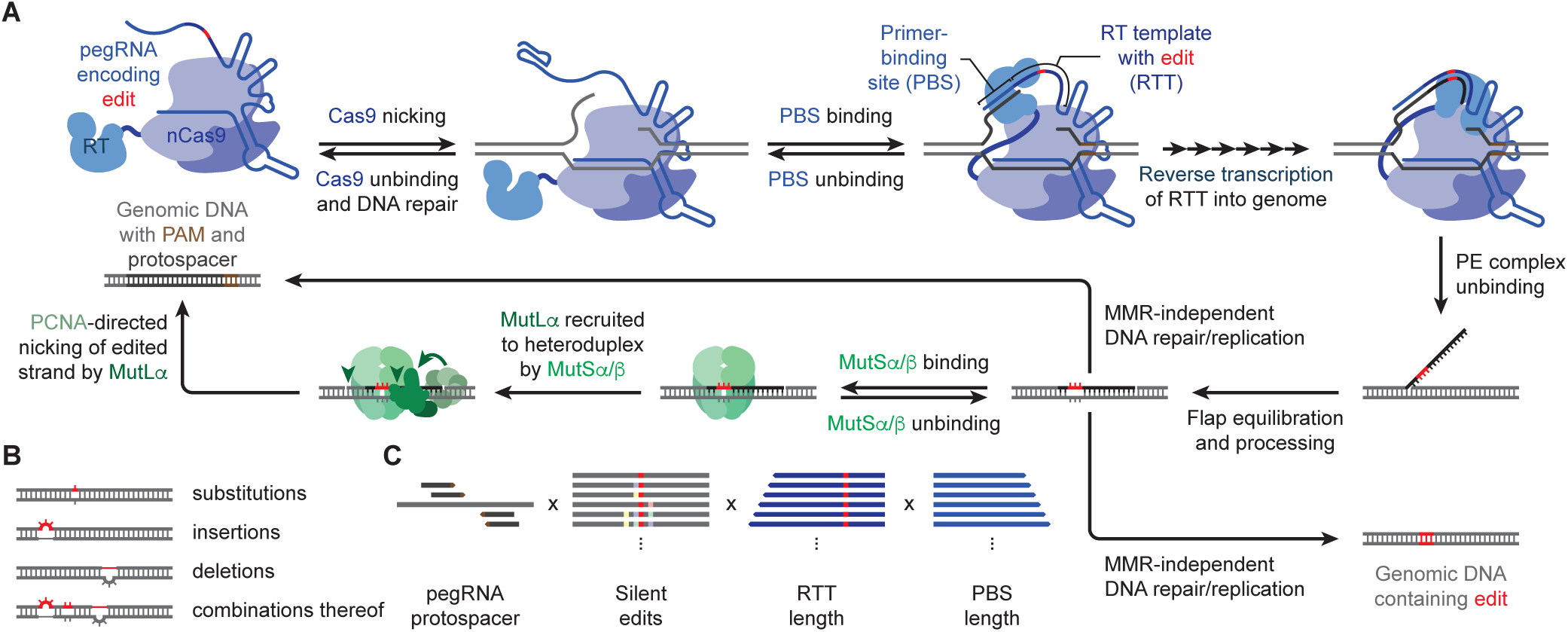
Design of prime editing (PE) strategies in a high-dimensional search space. **(A)** Prime editors write genetic information encoded by a prime editing guide RNA (pegRNA) directly into the genome of a cell with a reverse transcriptase (RT). After 3′-flap resolution, DNA repair processes, including mismatch repair (MMR), can resolve the heteroduplex into either the unedited sequence or the edited sequence. See also Figure S1. **(B)** Prime editors can effect small substitutions, insertions, deletions, and combinations thereof. **(C)** The search space of PE strategies is vast. Designing a pegRNA requires choosing (1) a PE protospacer, (2) MMR-evasive silent edits, (3) RT template (RTT) length, and (4) primer binding site (PBS) length.

Although the basic mechanistic steps of PE are well understood (Supplementary Figure 1), achieving efficient prime editing, especially in challenging cell types, can require evaluating hundreds or thousands of combinations of pegRNA design parameters in order to identify a pegRNA that maximizes PE efficiency^51–56^ (Figure 1C). Each pegRNA contains a spacer that directs the prime editor to its genomic target site, a reverse transcriptase template (RTT) that encodes the edit to be installed, and a primer-binding site (PBS) that hybridizes to the DNA primer to initiate reverse transcription. All three of these components are critical determinants of prime editing outcomes, and many options are often plausible for each. In addition, we and others previously showed that evading cellular mismatch repair (MMR) substantially increases PE efficiency, and therefore incorporating additional silent or benign mutations—of which there are typically many possibilities—can substantially improve PE outcomes by causing PE intermediates to be poor substrates for MMR^57,58^.

The vast number (typically, thousands) of potential pegRNAs that could support a given prime edit of interest, together with the high variability of PE efficiencies among these possible pegRNAs, is daunting for those who wish to use PE for both research and therapeutic purposes. To address this challenge, we and others have sought to develop a machine learning (ML) model that computationally identifies pegRNA designs with the highest potential to support efficient PE. The ability of such a model to decrease the number of experiments required to identify a high-performing PE strategy is dependent on the accuracy of the model. Currently published ML models for PE efficiency include PRIDICT^54,56^, DeepPrime^55,59^, and OPED^60^. Although these models achieve high accuracy on test datasets that were sampled from the large datasets on which they were trained, a substantial portion of each of these datasets consists of poor-performing pegRNAs that an experienced researcher would typically avoid during an initial screen. Consequently, although these models can reject low-efficiency pegRNAs in prospective design contexts, they struggle to accurately predict optimal pegRNAs amid a set of well-performing ones (*vide infra*). Moreover, these existing models are not based on the biological mechanism of PE. As a result, they may not take advantage of mechanistic insights that suggest key nodes that strongly determine PE outcomes, and their lack of correspondence with the molecular processes involved in PE makes it difficult to glean biological insights from their trained state and from their output.

Here, we describe the development of OptiPrime, an ML model that generates predictions of PE efficiency by directly incorporating knowledge about the biological mechanism of PE into its mathematical structure. By design, OptiPrime uses separate ML models to predict “pseudo-rates” of biochemical steps involved in the PE mechanism. Each pseudo-rate model only uses features that are biologically relevant to each process to make its prediction, ensuring a clean separation of tasks. The system of differential equations described by these rates is then integrated through time to obtain a final output prediction of editing efficiency. This approach contrasts with those used by PRIDICT^54,56^ and DeepPrime^55,59^, which incorporate all sequence features and representations directly into a single black-box predictor for PE efficiency. To train OptiPrime, we performed matched pegRNA–target site screens in both MMR-deficient HEK293T cells and MMR-competent HeLa cells, resulting in a dataset of 74,769 PE efficiencies across 1,290 unique target sites that enabled us to systematically study both MMR-dependent and MMR-independent sequence determinants of editing efficiency. While traditional ML models usually do not directly include different experimental conditions as covariates and instead rely on additional model fine-tuning with new data to make predictions in new contexts, the mechanistic structure of OptiPrime enabled its joint training on a dataset of 297,962 PE efficiencies collected across 40 experimental contexts in our laboratory and others. We demonstrate, through benchmarking and ablation studies, that OptiPrime offers best-in-class accuracy on PE efficiency prediction and that its performance is dependent on its mechanism-based mathematical structure.

Moreover, we show that the measurable effects of individual mechanistic steps in the PE process are consistent with their corresponding pseudo-rate constants in OptiPrime, that these pseudo-rates can be used to train a model of PE3 efficiency, and that OptiPrime can make useful predictions about twinPE (a dual-flap, two-pegRNA application of PE^61^), even though the model was never trained on twinPE data. Finally, we demonstrate the ability of OptiPrime to streamline the development of therapeutic PE applications in several prospective contexts, including the rapid development of a PE strategy to correct a pathogenic mutation in *Kif1a* that resulted in >40% *in vivo* prime editing efficiency in the bulk brain cortex of mice. Collectively, these findings demonstrate that OptiPrime augments the potential of PE by greatly accelerating the identification of efficient editing strategies, and that mechanistic ML models can provide insights into the dynamics of complex biomolecular processes, enabling modular, high-performance prediction of their outcomes in living systems.

## Results

### High-throughput prime editing library screens recapitulate structural and cellular determinants of mismatch repair

In a previous CRISPRi screen of DNA repair and DNA metabolism genes, we discovered that MMR is a major bottleneck for PE efficiency^57^. This insight led us to develop the PE4 and PE5 editing strategies, in which a dominant negative variant of the MMR protein MLH1 (MLH1dn) is co-delivered with the editing components to suppress MMR and favor other repair pathways that lead to desired PE outcomes^57^. Consistent with these results, we also found that the use of pegRNAs that install additional silent or benign mutations near the desired edit could substantially increase PE efficiencies by causing heteroduplex PE intermediates to contain sufficient mismatches between the edited and unedited strands to evade recognition by MMR proteins, which preferentially engage single-nucleotide or small mismatched regions^57^.

The MMR pathway is initiated upon mismatch binding by the MutSα or MutSβ complexes^62^, which together recognize small mismatches and small insertion–deletion loops^63,64^. To comprehensively evaluate the sequence determinants of binding by MutSα and MutSβ, we designed “Lib-MMR,” a set of 10,000 pegRNAs designed to target 200 randomly selected exonic sites in the human genome with diverse edit types, including single-base substitutions (1,592), contiguous substitutions of 2–5 bases (2,792), non-contiguous substitutions of 2–5 bases (2,780), deletions of ≤10 bases (1,000), insertions of ≤10 bases (1,000), protospacer-adjacent motif (PAM) edits with deletions (400), and PAM edits with insertions (400). Lib-MMR also included 36 positive control pegRNAs that we previously used to edit endogenous genomic sites (Table 1)^57^.

**Table 1.** Design of Lib-MMR.

| **Prime edit class** | **# members** |
|---|---|
| Single SNP | 1,592 |
| Contiguous SNPs | 2,792 |
| Non-contiguous SNPs | 2,780 |
| Deletions (<10 bp) | 1,000 |
| Insertions (<10 bp) | 1,000 |
| PAM SNP + deletion | 400 |
| PAM SNP + insertion | 400 |
| Endogenous control sites | 36 |
| Total | 10,000 |

Lib-MMR was designed to encompass a diverse set of prime edits in order to probe the sequence determinants of mammalian mismatch repair.

Although Lib-MMR contains diverse edit types and is therefore useful for assessing the determinants of binding by the MutSα/MutSβ complexes, we also sought to generate a dataset that would include many therapeutically relevant edits. Therefore, we designed “Lib-CV”, which comprises 10,406 pegRNAs that correct 944 known pathogenic variants in protein-coding regions from the ClinVar database^65^. In addition to a pegRNA that directly corrects each variant, we designed up to nine pegRNAs that also included benign “silent edits” that would not alter the corrected protein coding sequence (Table 2). Importantly, we designed the pegRNAs in Lib-MMR and Lib-CV with empirically determined heuristics to support efficient PE^52^ (Methods) so that an ML model that performs well on the resulting dataset will be required to distinguish between several candidates that perform well, rather than between rare candidates that perform well amidst many poor-performing alternatives.

**Table 2.** Design of Lib-CV.

| **Prime edit class** | **# members** |
|---|---|
| Correction to reference allele: |  |
| SNP reversion | 859 |
| Insertion reversion (deletion) | 6 |
| Deletion reversion (insertion) | 71 |
| Mixed reversion | 6 |
| Correction with silent edits: |  |
| SNP reversion | 8,600 |
| Insertion reversion (deletion) | 56 |
| Deletion reversion (insertion) | 723 |
| Mixed reversion | 62 |
| Total | 10,406 |

To assay PE efficiencies in a high-throughput manner, we packaged Lib-MMR and Lib-CV into lentiviral libraries of pegRNA–target site pairs. We transduced these libraries into cells at low multiplicity of infection (<0.3) to ensure that most transduced cells were only infected with a single lentivirus, selected for cells with integrated library members, then transfected these cells with PE construct plasmids to induce editing (Figure 2A). To thoroughly interrogate the effects of MMR on PE outcomes, we performed the screens with PE2 and PE4 in both HEK293T cells, which are partially MMR-deficient, and HeLa cells, which are MMR-proficient. As expected, we found that PE4 consistently outperformed PE2 across Lib-MMR. This effect was especially pronounced in HeLa cells, in which we observed a median 4.3-fold increase in editing rates with PE4 versus PE2, compared to only a median 1.5-fold increase in HEK293T cells (Figure 2B, 2C, Supplementary Figure 2A), consistent with previous observations that HEK293T cells are MMR-deficient. We chose to use the ratio of PE4 editing efficiency to PE2 editing efficiency (PE4:PE2) as a proxy for the propensity of a given edit to be reverted to the unedited sequence by MMR. We found that PE4:PE2 correlated more strongly with PE2 in HeLa cells (Spearman correlation ρ = −0.793) than in HEK293T cells (ρ = −0.390), suggesting that while MMR is the primary determinant of PE2 efficiency in MMR-proficient cell types, other factors determine PE2 efficiency in MMR-deficient cell types (Supplementary Figure 2B).

**Figure 2.**
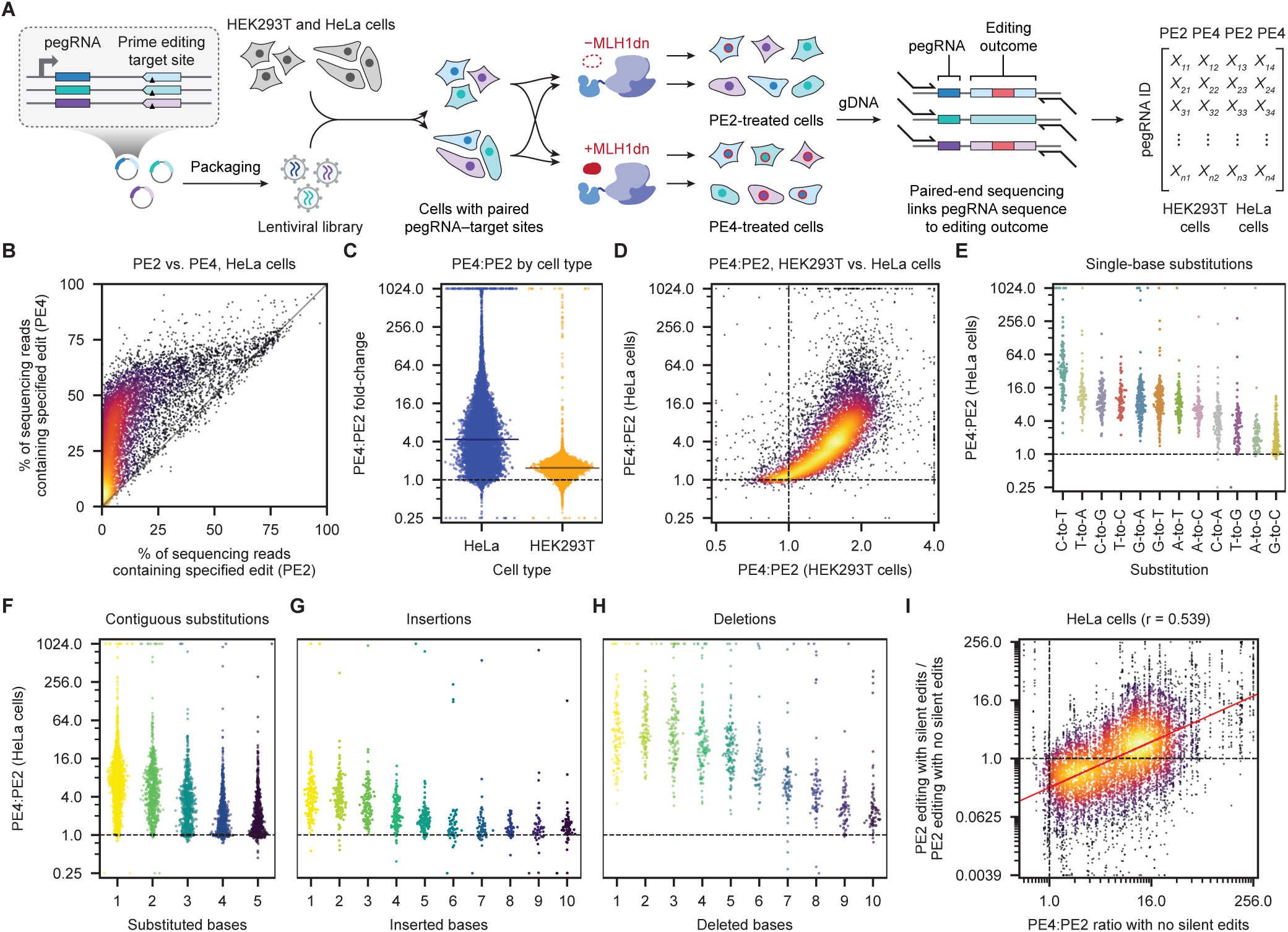
High-throughput lentiviral screens recapitulate the sequence determinants of mammalian mismatch repair (MMR). **(A)** Paired pegRNA–target site lentiviral libraries enable high-throughput evaluation of PE2 and PE4 outcomes in HEK293T and HeLa cells. **(B)** A scatter plot comparing editing efficiencies across Lib-MMR between PE2 (x-axis) and PE4 (y-axis) in HeLa cells. The gray line plots the equation y = x. PE4 consistently outperforms PE2 in HeLa cells. Brighter colors correspond to increased point density. See also Supplementary Figure 2A. **(C–H)** PE4:PE2 represents the ratio of PE4 editing efficiency to PE2 editing efficiency across library members in Lib-MMR. All PE4:PE2 values are plotted on a logarithmic scale. PE4:PE2 values below 0.25 (HeLa) or 0.5 (HEK293T) are clipped to each respective minimum value. PE4:PE2 values above 1024 (HeLa) or 4 (HEK293T) are clipped to each respective maximum value. **(C)** Beeswarm plots comparing PE4:PE2 between HeLa cells and HEK293T cells. PE4:PE2 is higher in HeLa cells than in HEK293T cells. **(D)** A scatter plot comparing PE4:PE2 across library members between cell types. PE4:PE2 correlates well between HEK293T and HeLa cells. Brighter colors correspond to increased point density. **(E)** Beeswarm plots of PE4:PE2 in HeLa cells for all twelve possible single-base substitutions, sorted by median PE4:PE2. See also Supplementary Figure 2E. **(F–H)** Beeswarm plots comparing PE4:PE2 in HeLa cells across varying lengths of (F) contiguous substitutions, (G) pure insertions, and (H) pure deletions. See also Supplementary Figure 2F–H. **(I)** A scatter plot comparing PE4:PE2 editing efficiency ratio of each edit to the ratio of PE2 editing efficiency with and without silent edits across Lib-CV in HeLa cells. Improvements in PE2 efficiency from including silent edits are positively correlated with PE4:PE2 efficiency ratios of the reference edit without any silent edits in HeLa cells. The red line is a total least squares regression line. Brighter colors correspond to increased point density.

We found that while PE4:PE2 values were generally lower in HEK293T cells than in HeLa cells as expected given the partial MMR deficiency of HEK293T cells, mean PE4:PE2 values correlated well between the cell types (ρ = 0.733), suggesting that trends in binding by MutSα and MutSβ are conserved (Figure 2D). Moreover, the PE4:PE2 ratios for single-base substitutions in Lib-MMR were highly consistent and linearly correlated with those obtained at endogenous genomic loci from our previous work (Pearson correlation *r* = 0.800, Figure S2C), with G-to-C and A-to-G edits resulting in the lowest median PE4:PE2 ratios, and C-to-T edits showing the highest median PE4:PE2 ratios^57^ (Figure 2E, Supplementary Figure 2D).

This ranking is consistent with the known propensity of the corresponding mismatched DNA intermediates to be repaired *in vitro* by HeLa cell extracts^66^. Although both A-to-G and C-to-T edits create mismatched T:G intermediates, these edits qualitatively show very different PE4:PE2 ratios, suggesting that T:G mismatches may be asymmetrically repaired by MMR or preferentially processed by an orthogonal repair pathway.

Additionally, we observed that edits consisting of longer contiguous substitutions resulted in decreasing PE4:PE2 values, with Spearman ρ values of –0.484 (HeLa) and – 0.427 (HEK293T) (Figure 2F, Supplementary Figure 2E). These data are consistent with structural observations that MutSα preferentially binds small mismatched regions rather than larger stretches of mismatched nucleotides^63^. Similarly, PE4:PE2 for both pure insertions and pure deletions decreased as the number of nucleotides inserted or removed increased, with deletion edits producing much higher PE4:PE2 than the corresponding insertion edits of the same length (Figure 2G, 2H, Supplementary Figure 2G, 2H). with strong negative rank-correlations for both pure insertions (HeLa ρ = −0.638; HEK293T ρ = −0.624) and pure deletions (HeLa ρ = −0.632; HEK293T ρ = −0.615). These data together suggest that known structural features of MMR proteins may underlie differences between PE2 and PE4 across a broad range of editing contexts.

In Lib-CV, we observed that including silent edits increased PE2 efficiency in an MMR-dependent fashion. The average log-fold-increase in PE2 efficiency at each site had positive, linear correlations with log-PE4:PE2 ratios of the simple reversion edit (HeLa *r* = 0.539; HEK293T *r* = 0.457, Figure 2I, Supplementary Figure 2I). Collectively, these data strongly reinforce our previous findings that including silent or benign mutations improves PE efficiency by shielding heteroduplex PE intermediates from engagement by cellular MMR machinery^57^. Moreover, by performing these screens in both HEK293T and HeLa cells, we confirmed that the bottleneck for efficient PE changes depending on the specific context in which the edit is performed, despite the common mechanism by which these edits are thought to occur.

### OptiPrime is a mechanism-based ML model of prime editing that enables high-performance prediction of prime editing efficiency and mismatch repair effects

A challenging aspect of modeling PE efficiency with ML is that any one of the many individual steps of the PE process can be a strong determinant of editing outcomes. For example, MMR on the unproductive strand results in the target site’s reversion to the original sequence, restarting the PE process from the beginning. However, a pegRNA that fails to engage the prime editor would not allow the target site to reach the heteroduplex state to be corrected by MMR. While it is theoretically possible for traditional ML models to learn the intricacies of these dynamics given sufficient training data, we hypothesized that an ML paradigm in which the model’s mathematical structure is based on a researcher-defined multi-step reaction mechanism would be particularly well suited for training a model of PE efficiency. Although no biophysical parameters are directly provided, we hypothesized that imposing this inductive bias on the structure of the model would allow for better model generalization through incorporation of domain knowledge. Moreover, this approach may allow relationships between key PE intermediates to be learned during training in ways that may accurately reflect effective rates between these intermediates in cells. Likewise, incorporating details regarding variations in experimental procedure is difficult with traditional ML models. While prior works on PE efficiency prediction have addressed this issue by tuning a separate model on each experimental context^55,56^, this solution narrows the applicability of the resulting models because it is unclear to what extent each individual model has learned rules intrinsic to PE itself that are general across many contexts.

Inspired in part by prior work in physics-informed neural networks, neural ordinary differential equations, and virtual whole-cell models^67–71^, we created OptiPrime, an ML model of PE efficiency based on our current understanding of the PE mechanism (Figure 3A, Supplementary Figure 1). In OptiPrime, a combination of deep neural networks and linear regression models are used to model the effective rates of steps in our hypothesized mechanism of PE (“pseudo-rates”). These rates form a system of ordinary differential equations that can be integrated through time to evaluate the editing at the terminal timepoint for a given editing experiment (Figure 3B–C). While it would be experimentally impractical to create an ML training dataset with direct measurements of ground-truth rates in cells for many prime edits, we reasoned that pseudo-rates predicted using only handpicked salient features would represent their respective distinct steps in the PE mechanism. For a detailed discussion of the OptiPrime architecture and the considerations made in its design, see Supplementary Text 1.

**Figure 3.**
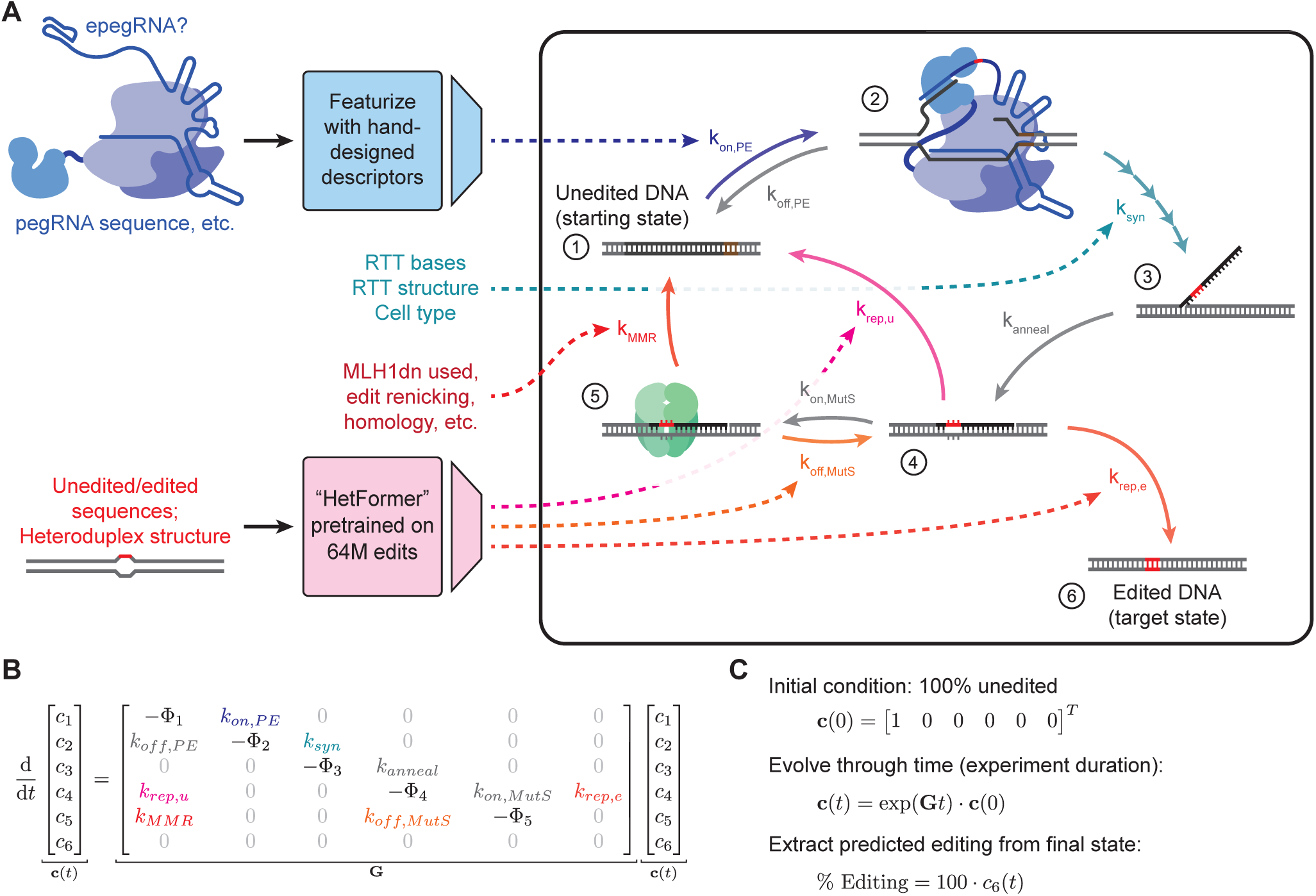
OptiPrime is a mechanism-based machine learning model of prime editing efficiency. **(A)** The mechanism of prime editing used by OptiPrime as an inductive bias. Numbers (1–6) in circles represent the different genomic states that are used in the OptiPrime mechanism. Colored rates represent those which are predicted by machine learning models (left), while gray rates represent those that are kept constant (for a given cell type). **(B)** The differential equation that governs how the concentration of each state (c_i_) evolves over time. Predicted and constant rates are placed in the generator matrix at off-diagonal positions. −Φ_i_ terms are the negative sum of all rates in the row, which maintains mass balance. **(C)** An overview of the procedure used to generate predictions with OptiPrime.

In brief, we divided our model into mechanistic steps that are mediated by the prime editor or those that only involve DNA repair machinery (Figure 3A). We modeled the former with hand-crafted features that were selected for their relevance to either DNA binding and nicking (k_on,PE_) or flap synthesis (k_syn_). We then explicitly modeled the MMR pathway by including a reversible step for heteroduplex binding by the MutSα/MutSβ complexes. We represented MMR-independent heteroduplex repair as well as the off-rate of the MutSα/MutSβ complexes with the “HetFormer”, a neural network architecture we developed for this work inspired by the EvoFormer used by AlphaFold^72,73^. We pre-trained HetFormer on 64 million simulated heteroduplexes (Supplementary Figure 3, Supplementary Text 1). We trained OptiPrime on a dataset of 297,962 PE experiments that combines the data from our paired pegRNA–target site screens with prime editing data collected by other laboratories in previous studies^54–56^ to maximize the likelihood that the resulting model learns aspects of the PE mechanism that are general across all tested conditions.

To estimate model performances on unseen data, we split our dataset five ways, stratified by protospacer sequence (to avoid information leakage across splits). For each split, we trained models on four of the data partitions and evaluated on the other partition (5-fold cross-validation). Under these conditions, we found that OptiPrime consistently achieved high predictive performance on all training datasets, including those collected in this study (mean *r* = 0.693, ρ = 0.745, Figure 4A). We also compared OptiPrime to the previously reported DeepPrime^55^ and PRIDICT^54,56^ models (Figure 4A). We found that OptiPrime obtained significantly higher cross-validation rank correlations than DeepPrime-FT (mean *r* = 0.325, ρ = 0.399), PRIDICT2.0-HEK (mean *r* = 0.426, ρ = 0.538), and PRIDICT2.0-K562 (mean *r* = 0.516, ρ = 0.590) on the datasets collected in this work (all Steiger’s *p* < 10^-323^, Figure 4A).

**Figure 4.**
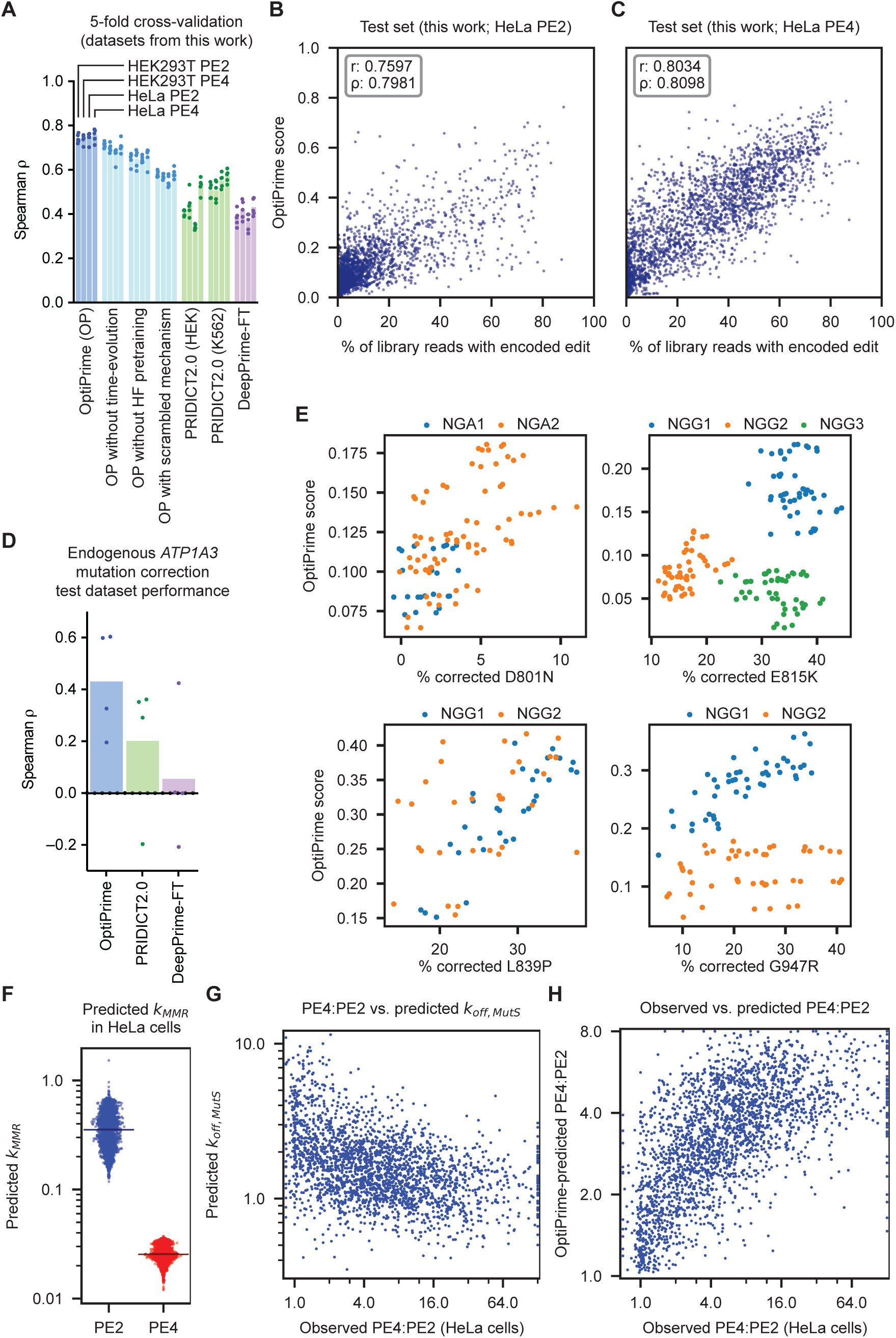
OptiPrime accurately predicts prime editing efficiencies across diverse experimental contexts. **(A)** Spearman ρ values on different 5-fold cross-validation splits for the data collected in this study. The results for the final version of OptiPrime are in blue, those for ablation studies are shown in light blue, those for the PRIDICT2.0 models are shown in green, and those for the DeepPrime models are shown in purple. “OP without time-evolution” refers to replacing the mechanistic time-evolution layer in OptiPrime with a linear model that combines the outputs of the individual rate models. “OP without HF pretraining” refers to training OptiPrime with randomly initialized HetFormer weights, rather than from weights obtained from pretraining on 64,000,000 synthetically generated edits. “OP with scrambled mechanism” refers to training a version of OptiPrime in which the ML model that predicts the rate of each mechanistic step has been assigned to steps randomly, rather than based on the mechanism of prime editing. “PRIDICT2.0 (HEK)” and “PRIDICT2.0 (K562)” refer to the latest publicly available models trained on each cell type, respectively. DeepPrime-FT uses the HEK293T PE2max-e, HEK293T PE4max-e, HeLa PE2max, and HEK293T PE4max-e (since there is no DeepPrime-FT version for HeLa PE4max-e) models to predict values for HEK293T PE2max, HEK293T PE4max, HeLa PE2max, and HeLa PE4max, respectively. **(B, C)** Scatter plot of test set predictions with OptiPrime for the data collected in this study in HeLa cells with (B) PE2 and (C) PE4. **(D)** Comparison of rank correlations between OptiPrime, PRIDICT2.0, and DeepPrime-FT on data collected at the endogenous ATP1A3 locus for correcting four pathogenic mutations. **(E)** Scatter plots of ATP1A3 correction with OptiPrime scores. Colors represent the spacers, as denoted in ref. ^47^. **(F)** Beeswarm plots of predicted MMR in HeLa cells across the test sets collected in this work. As expected, predicted k_MMR_ decreases when changing from a PE2 context to a PE4 context. **(G)** A scatter plot of observed PE4:PE2 editing efficiency ratio (PE4:PE2) in HeLa cells vs. predicted k_off,MutS_ across the test sets collected in this work. As expected, decreased k_off,MutS_ corresponds to an increased binding affinity for the MutSα/MutSβ complexes, resulting in increased MMR correction (as quantified by PE4:PE2). **(H)** A scatter plot comparing observed PE4:PE2 in HeLa cells to the ratio of the corresponding predictions by OptiPrime. Predicted PE4:PE2 correlates well (r = 0.582, ρ = 0.668), indicating that OptiPrime has learned the sequence determinants of MMR correction propensity.

To determine the design features that contribute to OptiPrime’s predictive performance, we also performed an ablation study, in which several versions of OptiPrime were re-trained with each component systematically removed. Overall, we found that the results achieved by OptiPrime were dependent on all of its key design features—removing any component resulted in significantly worse cross-validation performance on all of our datasets, including the mechanistic time-evolution layer and the pretraining of the HetFormer (all Steiger’s *p* ≤ 2×10^-12^, Figure 4A). Moreover, “scrambling” the reactions by re-assigning the rate models of the mechanistic graph to randomly chosen reaction edges also significantly deteriorated the model performance (all Steiger’s *p* < 10^-323^, Figure 4A), indicating that OptiPrime’s high performance is derived from the correspondence between its architecture and the underlying PE mechanism. Collectively, these findings indicate that the above design choices each contribute to OptiPrime’s high accuracy.

With these cross-validation results in hand, we computed predictions with the final version of OptiPrime on held-out test sets sampled from each dataset (Figures 4B, 4C, Supplementary Figure 4). The cross-validation results translated well to the test set across all four experimental conditions tested in this work (mean *r* = 0.723, ρ = 0.775). Lastly, we compared OptiPrime’s predictions to those generated by PRIDICT2.0 and DeepPrime on several test sets of prime edits from arrayed experiments that we performed during the course of our work on correcting mutations in *ATP1A3* that cause alternating hemiplegia of childhood^47^ (Figure 4D, 4E). We observed high correlations between OptiPrime and empirical editing for D801N (ρ = 0.604) and L839P (ρ = 0.599) and modest correlations for E815K (ρ = 0.326) and G947R (ρ = 0.196). We noted that these relatively poor performances could be primarily attributed to bad predictions with protospacers E815K-NGG3 and G947R-NGG2, which both supported efficient PE in spite of their poor predicted protospacer scores by the CRISPR nuclease sgRNA model Rule Set 3^74^. Indeed, removing E815K-NGG3 and G947R-NGG2 improved OptiPrime’s rank correlation on the respective test sets to ρ = 0.786 and ρ = 0.787, respectively (Figure 4E). In comparison, neither PRIDICT2.0 nor DeepPrime-FT were able to achieve rank-correlations above 0.43 on any of the four test sets, and both models generated predictions with negative rank-correlations at one site, suggesting that following the predictions of PRIDICT2.0 or DeepPrime-FT can sometimes produce results that would be on average results worse than selecting at random from the candidate pegRNAs we evaluated. Taken together, these results show that OptiPrime models PE efficiency with an accuracy that can exceed that of previously published models.

Another potential benefit of OptiPrime’s mechanistic architecture is the possibility that the resulting learned pseudo-rates may provide insights into specific steps of the PE process. Unlike non-mechanistic ML models, in which outcomes cannot be disentangled from their influence by individual model features, the mechanistic nature of OptiPrime enabled us to investigate in greater detail how input features influence each pseudo-rate. For example, we found that predicted *k*_MMR_, which takes MLH1dn use as an input, decreases by a median of 14-fold under PE4 conditions compared to under PE2 conditions, consistent with our understanding that the use of MLH1dn in PE4 inhibits MMR (Figure 4F).

To further probe OptiPrime’s ability to evaluate the role of MMR across diverse edits, we investigated the relationships between PE4:PE2 and the three pseudo-rates predicted by HetFormer: *k*_rep,u_, *k_r_*_ep,e_, and *k*_off,MutS_. In our proposed PE mechanism, we consider binding of MutSα/MutSβ complexes to an editing intermediate to be necessary for that edit to be reverted to the unedited sequence by MMR machinery. In OptiPrime, we assume that the on-rate is governed by diffusion and that all variation in binding affinity can be attributed to the off-rate. We found a significant negative rank-correlation between predicted *k*_off,MutS_ rates and empirical PE4:PE2 (HeLa ρ = −0.42, *p* = 1.5×10^-104^) across our test set, consistent with our hypothesis that tighter binding by the MutSα/MutSβ complexes (i.e., a lower off-rate) results in increased reversion by MMR (i.e., increased PE4:PE2) (Figure 4G). Moreover, the ratio of predicted PE4 efficiencies to predicted PE2 efficiencies with OptiPrime correlates well with observed PE4:PE2 values (*r* = 0.582, ρ = 0.668), indicating that the HetFormer has learned the sequence determinants salient to MMR in the context of PE (Figure 4H) and that OptiPrime is well-suited for selecting MMR-evading silent edits. We speculate that this ability arises from the fact that OptiPrime was jointly trained on both PE2 and PE4 simultaneously, rather than through fine-tuning on PE4 data.

### Pseudo-rates learned by OptiPrime enable PE3 and twinPE predictions

We next asked whether OptiPrime pseudo-rates could be used to predict outcomes of PE variants such as adding a nicking guide RNA (‘PE3’^49^) or using two pegRNAs to generate complementary edited strands (‘TwinPE’^61^), despite a lack of training data on these variants. In PE3, a nicking sgRNA (nsgRNA) is used to drive DNA repair towards incorporation of the edit by nicking the DNA strand complementary to the PE protospacer (Figure 5A). Analogous to the relationship between PE2 and PE4, we previously found that including MLH1dn could improve PE3 efficiencies and reduce indel formation, a strategy we designated PE5^57^. We assembled a dataset of 940 PE3 (or PE5) efficiencies paired with their corresponding PE2 (or PE4) efficiencies with 159 prime edits collected at endogenous sites in human cells. Although some PE strategy design tools allow users to nominate nsgRNAs for PE3, these tools give recommendations solely on the basis of predicted Cas9 nuclease efficiencies for indel generation^51,54,56^. While creating double-strand DNA breaks by Cas9 nuclease is a prerequisite for indel formation in these contexts, indel sequence outcomes are shaped by various end-joining pathways (i.e., c-NHEJ and MMEJ). This reasoning is consistent with our observation that neither DeepSpCas9 (mean ρ = −0.086, all *p* > 0.05)^75^ nor Rule Set 3 (mean ρ = −0.084, all *p* > 0.05)^74^ scores provided meaningful predictive PE3 editing efficiency signal on any test dataset (Figure 5B), although we note a modest correlation between DeepSpCas9 (mean ρ = 0.241) or Rule Set 3 (mean ρ = 0.221) scores and PE3 indel efficiencies. These data suggest that nuclease scores are not useful for predicting optimal nsgRNAs in the context of PE3, motivating the development of bespoke models of PE3 efficiency.

**Figure 5.**
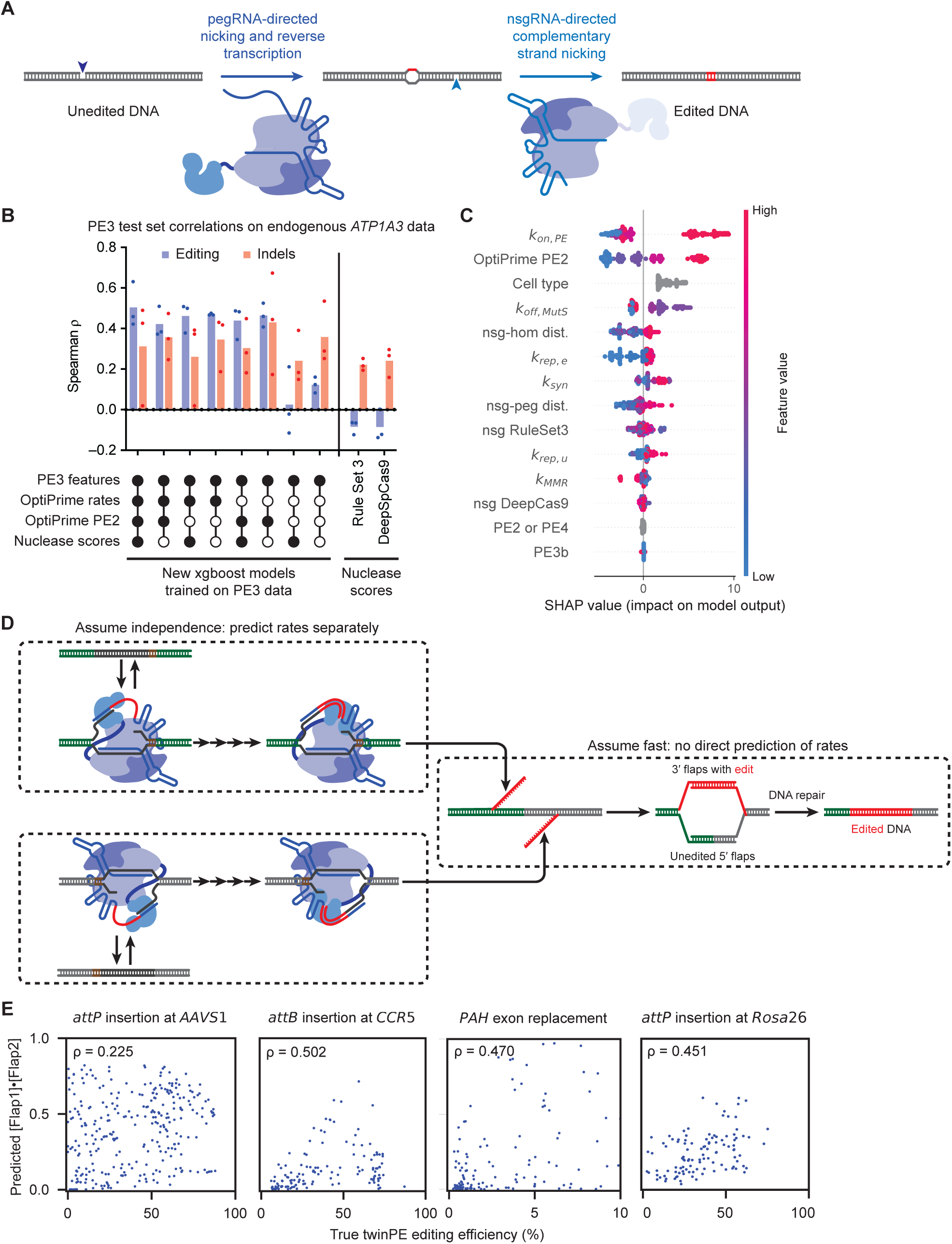
OptiPrime rates enable prediction of PE3 and twinPE outcomes. **(A)** In PE3, a complementary strand-nicking sgRNA (nsgRNA) drives editing towards the desired product. **(B)** Test set correlations of PE3 editing data taken from our work correcting three pathogenic ATP1A3 mutations^47^. **(C)** Beeswarm plots of SHAP values for PE3 editing predictions on test sets. **(D)** TwinPE installs longer edits by synthesizing complementary flaps with two pegRNAs that target complementary strands. We predict twinPE efficiencies with OptiPrime by assuming that 3’-flap synthesis is rate-limiting, and that synthesis rates of each flap is independent of the other. **(E)** Scatter plots showing zero-shot prediction correlations of twinPE efficiencies on four datasets of twinPE efficiencies taken from our previous work^61,77^.

Transfer learning is a ML approach in which a model trained on one task is repurposed for a different, typically related, task. A common strategy for transfer learning is to fix the base model’s weights and use its outputs or computational intermediates as model inputs for the new task. Although the detailed mechanisms by which PE3 improves editing efficiency are not fully understood, we hypothesized that the pseudo-rates learned by OptiPrime could serve as mechanistically relevant features for predicting PE3 efficiencies. We trained gradient-boosted tree models to predict PE3 editing efficiencies and indel frequencies based on these pseudo-rates, predicted OptiPrime PE2 (or PE4) efficiencies, as well as other hand-extracted features (Methods) and found that the resulting model was able to achieve good predictive correlation for each PE3 test dataset collected during the course of prime editing experiments correcting pathogenic mutations in *ATP1A3* (mean ρ = 0.534, Figure 5B).

Consistent with our observation that nuclease efficiency scores alone do not predict PE3 nsgRNA efficiency, we found that removing OptiPrime-predicted pseudo-rates and editing efficiencies as input features abrogated predictive power on our test sets (mean ρ = 0.025, all *p* > 0.05, Figure 5B). Consistent with these findings, when we quantified the contribution of each input feature to our PE3 model’s test set prediction performance with the widely used SHAP explanation technique^76^, we found that features taken from OptiPrime were among the most important factors influencing its outcomes—*k*_on,PE_, OptiPrime-predicted editing efficiency, and *k*_off,MutS_ were all among the top five predictive features (Figure 5C). We also found that including only the final predicted PE2 (or PE4) efficiency from OptiPrime could still achieve some predictive power, albeit lower than when pseudo-rates are included (mean ρ = 0.464, Figure 5C). Taken together, these results further indicate that integrating inferred mechanistic dynamics into models of PE efficiency with nicking guide RNAs (PE3) is crucial for accurate performance.

We also hypothesized that OptiPrime could predict the outcomes of twinPE, in which two pegRNAs targeting complementary genomic DNA strands are used to generate two 3′ flaps that anneal to each other, obviating the need for second-strand DNA synthesis to generate stable PE products (Figure 5D). While a traditional ML model trained on PE data would be unsuitable for such a task because predictions about the relevant flap synthesis steps cannot be disentangled from those for irrelevant DNA repair steps, mechanistic ML could enable us to rewire pseudo-rate models to generalize to processes such as twinPE with different but related mechanisms compared to those used to generate the base model.

Although it is not possible to model DNA repair in twinPE using OptiPrime because twinPE does not create an unresolved heteroduplex intermediate as traditional PE does, we assumed that 3′-flap installation efficiency is the rate-limiting step in the twinPE mechanism. We thus chose to predict twinPE efficiencies by taking 3′-flap synthesis pseudo-rates from OptiPrime for each individual pegRNA, ignoring the pseudo-rates for all DNA repair steps downstream of flap synthesis (Figure 5D). We collected a dataset of 667 twinPE outcomes across four endogenous loci from our previous reports disclosing twinPE^61^ and eePASSIGE^77^, a technique that uses PE to insert landing sites for gene integration with the Bxb1 recombinase. We found that this procedure achieves moderate predictive performance across the four twinPE datasets (mean ρ = 0.412, Figure 5E) despite never having been trained on any twinPE data, demonstrating that OptiPrime has learned features inherent to the steps in the PE mechanism that can then be disentangled and used separately for new mechanisms. Collectively, these data demonstrate that mechanistic ML models can facilitate downstream fine-tuning or rewiring to generate predictions on new, mechanistically related processes.

### OptiPrime accelerates the development of therapeutic prime editing strategies

To establish the utility of OptiPrime in finding efficient PE strategies in various therapeutic contexts, we designed pegRNAs capable of effecting several therapeutically relevant edits, including correction of four pathogenic mutations in the *CFTR* gene and one pathogenic mutation in the *COL7A1* gene, orthogonalization of the IL2 receptor, and correction of a pathogenic *Kif1a* mutation in a mouse model of KIF1A-associated neurological disorder (KAND) (Methods). We tested these pegRNAs in various cell types, including in human primary T cells and in embryonic fibroblasts from the *Kif1a*^lgdg^ model mouse. Finally, we used the results from an OptiPrime-streamlined optimization campaign in an *in vivo* study to demonstrate how OptiPrime can greatly accelerate the development of PE strategies to correct pathogenic mutations that cause rare diseases.

Cystic fibrosis (CF) is caused by mutations in the CF transmembrane regulator (*CFTR*) gene that encodes the CFTR protein, an anion transporter of Cl^-^ and HCO_3_^-82^. We recently reported the development of a PE strategy for correcting CFTR p.F508del, a mutation found in 85% of people with CF^12^. Our efforts towards developing this strategy involved several cycles of iterative pegRNA optimization over the course of several years, resulting in the identification of “SE2”, an epegRNA encoding several silent edits that corrects p.F508del with 11% efficiency in HEK293T cell lines homozygous for *CFTR* p.F508del when used in a PE2 strategy (without a nicking guide RNA). Assaying the eight designs with the highest OptiPrime-predicted editing efficiencies enabled us to identify an epegRNA capable of correcting this mutation with 22% efficiency in a PE2 strategy, a 2.0-fold increase over that achieved by SE2 (Figure 6A, Supplementary Figure 5A). In contrast, none of the 16 top-scoring designs proposed by either PRIDICT2.0 or DeepPrime was able to achieve over 1% editing (Figure 6A).

**Figure 6.**
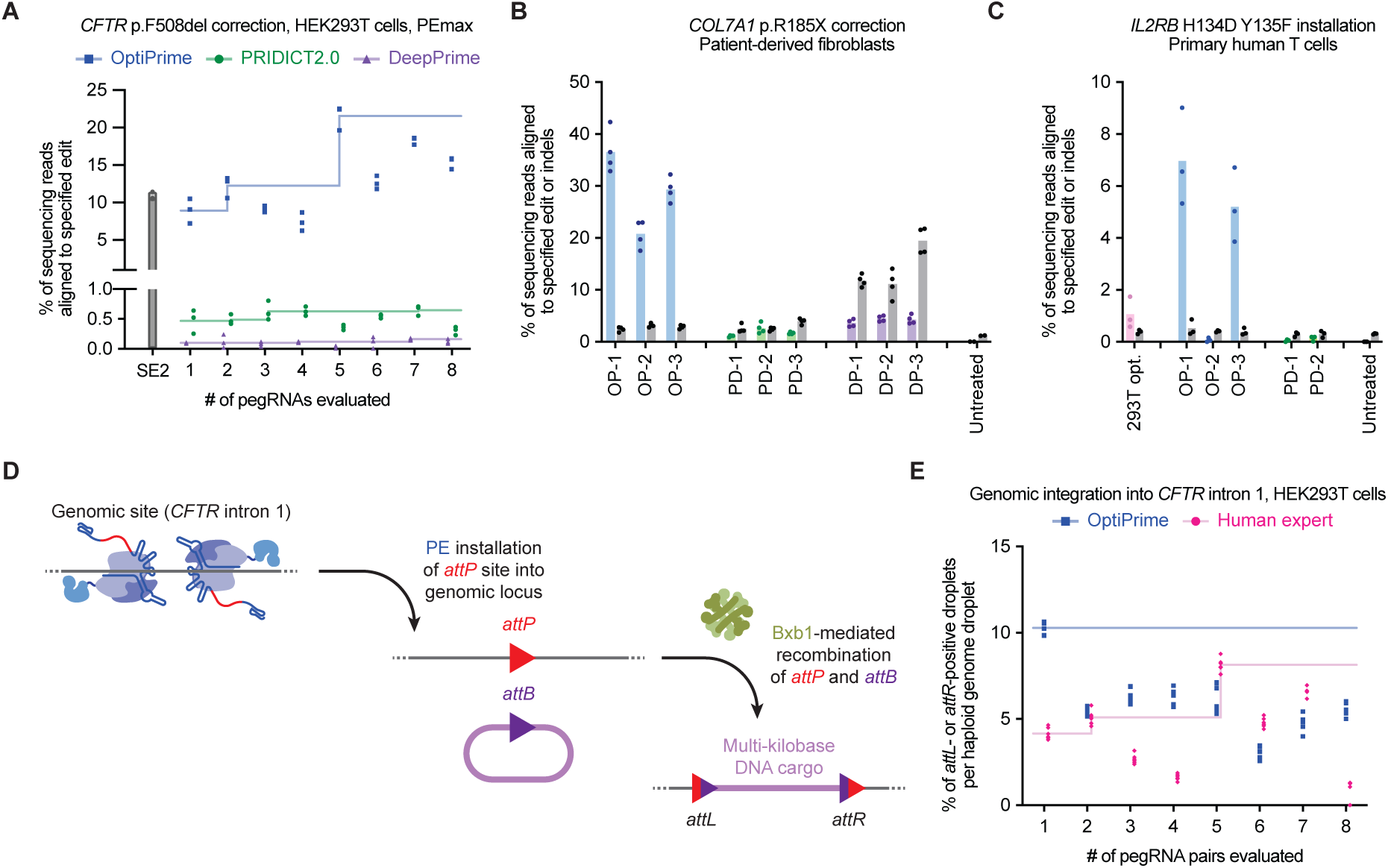
OptiPrime streamlines identification of efficient therapeutic prime editing strategies in vitro. **(A)** Correction of the *CFTR* p.F508del mutation with OptiPrime, PRIDICT2.0, and DeepPrime in HEK293T cells. Points denote individual experimental editing efficiencies (N = 3) for each ranked pegRNA on the x-axis, while lines indicate the maximum editing efficiency achieved after evaluating the top x-ranked pegRNAs. **(B)** Correction of the *COL7A1* p.R185X mutation with OptiPrime, PRIDICT2.0, and DeepPrime in fibroblasts derived from patients with recessive dystrophic epidermolysis bullosa. **(C)** Installation of the orthogonal IL2 receptor (IL2RB H134D Y135F) in primary human T cells. “293T opt.” denotes editing efficiency with a previously reported pegRNA without silent edits optimized by hand in HEK293T cells^57^. **(D)** PASSIGE is a strategy for the insertion of large DNA cargo into mammalian cells^61,77^. First, PE is used to install an attP site into a genomic locus. Then, a DNA donor containing the corresponding attB site is recombined into the genome with a large serine recombinase, such as Bxb1. **(E)** Installation of a gene-sized cargo into intron 1 of *CFTR* in HEK293T cells with eePASSIGE with either OptiPrime-nominated pegRNA pairs or expert-designed pegRNA pairs. Points denote individual experimental editing efficiencies (N = 3) for each ranked pegRNA pair on the x-axis, while lines indicate the maximum editing efficiency achieved after evaluating the top x-ranked pegRNA pairs.

Although the CFTR p.F508del mutation is present in the majority of people with CF, more than 2,000 *CFTR* variants have been identified in the human population, over 700 of which are verified to cause CF^82^. We used OptiPrime to score and select pegRNAs that are capable of correcting either the p.W1282X or p.G542X nonsense mutations, which are not responsive to small-molecule CFTR modulators such as Trikafta. For both mutations, assaying the eight top OptiPrime-predicted epegRNAs resulted in higher editing than assaying the same number of epegRNAs scored by PRIDICT2.0 or DeepPrime (Supplementary Figure 5B–E). We noted that, despite allowing OptiPrime to evaluate silent edits proximal to p.W1282X, the optimal strategy proposed by OptiPrime did not have any silent edits, indicating that the higher editing efficiency of the OptiPrime-nominated design is due to an RTT+PBS length combination that was more optimal than those nominated by PRIDICT2.0 or DeepPrime, rather than an advantage conferred by silent edit designs. Moreover, the PE-suitable NGG PAM closest to CFTR p.G542X would place the corrective T-to-G edit 45 bp downstream of the PE nick. Although OptiPrime, PRIDICT2.0, and DeepPrime do not allow design of edits this distance away, OptiPrime alone was able to identify a protospacer that places the c.1624G>T mutation 4 bp downstream from the PE nick with an NGA PAM that is compatible with a Cas9-VRQR variant^83^. For the sake of comparison, we assessed the RTT+PBS length combinations output by PRIDICT2.0 and DeepPrime by replacing the NGA PAM used by the OptiPrime-selected protospacer with NGG.

Next, we tested how the pegRNAs from OptiPrime performed in primary human cell types, which were not used to generate any of OptiPrime’s training data. Recessive dystrophic epidermolysis bullosa (RDEB) is caused by mutations in the *COL7A1* gene that result in impaired function of type VII collagen, leading to breakdown of the dermal–epidermal junction and a severe blistering phenotype^84^. Using OptiPrime, PRIDICT2.0, and DeepPrime, we chemically synthesized the top three epegRNAs as evaluated by each model and electroporated them into RDEB patient-derived fibroblasts with *in vitro*-transcribed mRNA encoding PEmax. We found that the best of the three OptiPrime-selected pegRNAs achieved 37% correction, while none of the six pegRNAs selected by PRIDICT2.0 and DeepPrime were able to effect above 5% correction (Figure 6B). We note that the DeepPrime-nominated pegRNAs showed high levels of indels, which we speculate may arise from its selection of pegRNAs with short RTT homology lengths. Notably, the levels of editing from OptiPrime-generated pegRNAs were also higher than those we previously achieved with adenine base editing^16^, demonstrating that OptiPrime is capable of designing PE strategies as efficient as those using base editing in difficult cell types without additional optimization.

We also chose to revisit the installation of the *IL2RB* H134D Y135F variant that enables selective stimulation of T cells with a corresponding orthogonal IL-2 cytokine^85^. We electroporated primary human T cells with PEmax-encoding mRNA and chemically synthesized pegRNAs. In this assay, we compared three OptiPrime designs that include silent edits to three designs without silent edits, two of which were designed with PRIDICT2.0 (which does not generate silent edits) and one of which we previously identified as the top performer from a manual assessment of 24 pegRNAs in HEK293T cells^57^. We did not include any comparisons with DeepPrime, since it is limited to making ≤3-bp changes. We found that the best of the three OptiPrime-designed pegRNAs achieved 7.0% editing in a PE2 context with synthetic pegRNAs. In comparison, the best pegRNA without any silent edits, which was optimized beforehand in HEK293T cells, achieved only 1.1% editing, and both of the PRIDICT2.0-generated pegRNAs yielded ≤0.1% editing (Figure 6C). Taken together, these results demonstrate that the silent edit combinations identified by OptiPrime can improve editing efficiency at previously challenging targets and in cell types beyond those used to generate OptiPrime’s training data, including primary human T cells.

Given OptiPrime’s ability to predict twinPE efficiencies, we designed twinPE epegRNAs capable of installing the attP landing site for Bxb1 recombinase in intron 1 of *CFTR*, as one step towards a potential mutation-agnostic therapy for people with CF caused by rare mutations (Figure 6D). We performed eePASSIGE^77^ integration experiments comparing one-pot integration efficiencies of eight epegRNA pairs nominated by OptiPrime against eight epegRNA pairs designed by a human PE expert with assistance with CRISPick^74,86,87^. We found that the top OptiPrime-nominated epegRNA pair yielded 10% integration efficiency of a multi-kilobase plasmid cargo, compared to 8.0% achieved by the best-performing human expert-designed epegRNA pair (Figure 6E).

Finally, we tested if OptiPrime could streamline efforts to address a longstanding need in medicine—the development of *N*=1 treatments for *de novo* genetic mutations that result in disease^88^. While high-throughput sequencing allows us to diagnose the genetic causes of some diseases rapidly, the development of a corresponding genome editing treatment has until recently been too slow to rescue patient phenotypes for most urgent genetic diseases. In a study led by Musunuru, Ahrens-Nicklas, and coworkers, we recently reported the development of the first personalized gene-editing treatment of a rare genetic disease, in which an infant was dosed with a bespoke base editor developed specifically to correct the infant’s unique mutation in *CPS1* at only seven months of age^48^. Developing this treatment so rapidly was possible due to the relatively small design space of base editing strategies; such a timeline would be more difficult for PE, given the far larger design space of pegRNAs.

We envision a pipeline in which detection of a pathogenic mutation is followed by streamlined generation of a small number of promising candidate corrective pegRNAs with OptiPrime. These candidates are then tested directly in patient-derived cells to rapidly identify efficient editing strategies (Figure 7A). As a proof-of-concept for this strategy, we used OptiPrime to design pegRNAs to correct the “leg dragger” mutation in *Kif1a*^lgdg^ mice. This spontaneous missense mutation results in the Kif1a p.L181F protein change and causes spastic paraplegia in homozygous *Kif1a*^lgdg^ similar to the defects observed in patients with pathogenic mutations in *KIF1A*^89,90^.

**Figure 7.**
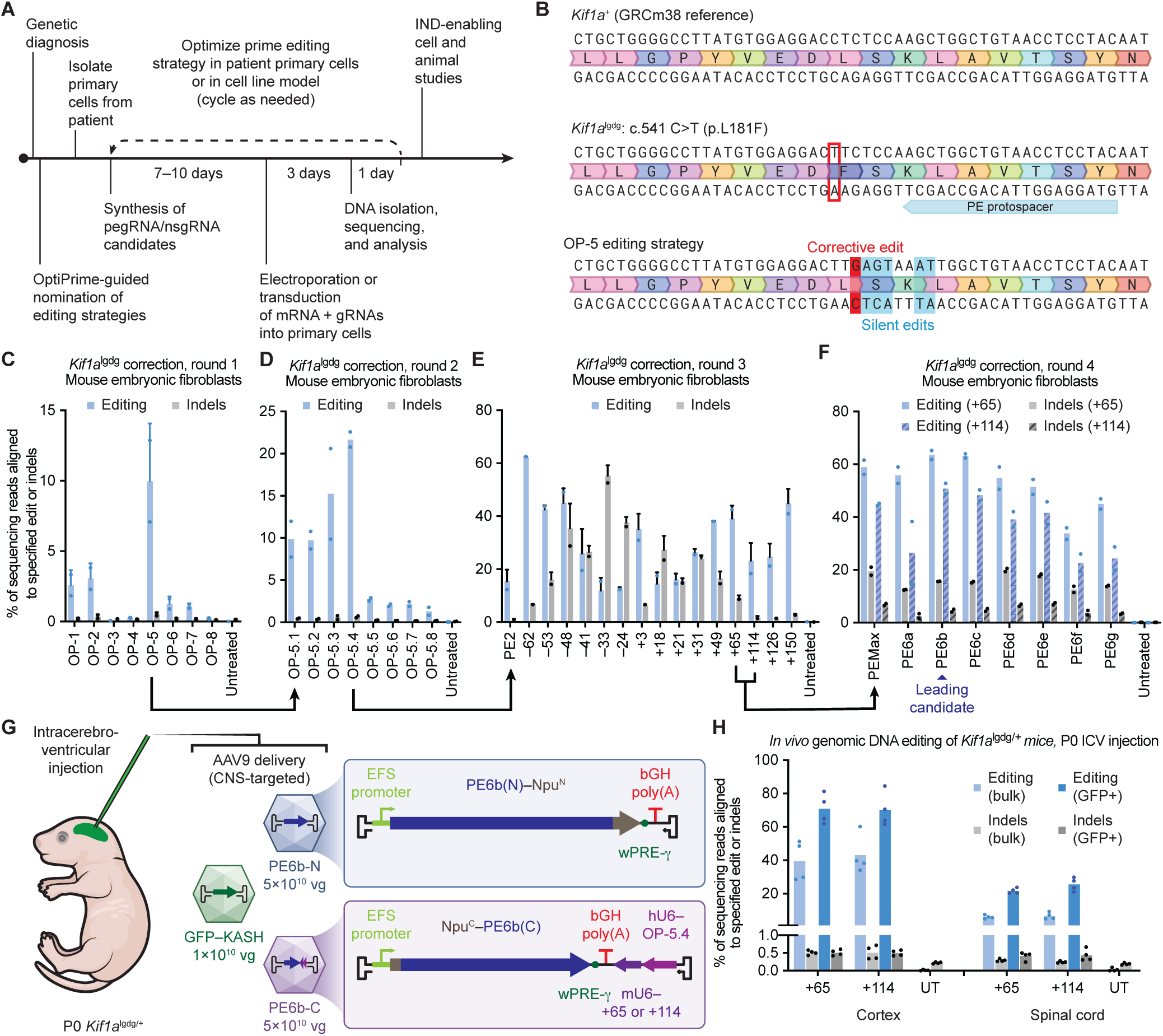
OptiPrime accelerates the development of corrective strategies for pathogenic mutations in vivo. **(A)** An experimental outline for rapid development of therapeutic prime editing strategies with OptiPrime. Rather than generating a cell line bearing the mutation of interest, we propose directly editing patient-derived cells through electroporation of in vitro-transcribed mRNA and synthetic pegRNA/nsgRNA constructs or viral transduction. **(B)** *Kif1a* genotypes relevant to this study. *Kif1a*^+^ refers to the genotype from the GRCm38 reference assembly. *Kif1a*^lgdg^ is caused by a C-to-T transition mutation resulting in the p.L181F protein change. The editing strategy used by the OP-5 family of pegRNAs changes a different codon position than the pathogenic mutation and includes five non-coding silent edits. **(C–F)** Lead optimization of a corrective strategy for *Kif1a*^lgdg/+^ in mouse embryonic fibroblasts through electroporation of PE mRNA, synthetic pegRNAs, and synthetic nsgRNAs. **(C)** OP-5 was identified in an initial assessment of eight pegRNAs encoding different silent edit strategies. **(D)** OP-5.1 was identified in a follow-up experiment investigating eight RTT+PBS lengths for OP-5. **(E)** Assaying 15 nsgRNAs identified the +65 and +114 nsgRNA sequences as promising candidates for further assessment. **(F)** Assessing engineered and evolved PE protein sequences identified PE6b as our lead candidate. **(G)** In vivo correction strategy of *Kif1a*^lgdg^. Dual AAV9 encoding split PE6b, the in vitro-optimized pegRNA (converted to an epegRNA), and the in vitro-optimized nsgRNA were delivered through intracerebroventricular injection into P0 *Kif1a*^lgdg/+^ mice. **(H)** In vivo editing of *Kif1a* genomic DNA in mouse cortex and spinal cord. Genomic DNA from both bulk mouse tissue and GFP-positive nuclei (to enrich for transduced cells) was evaluated for PE efficiency 4 weeks post-injection.

We designed pegRNAs to correct the *Kif1a*^lgdg^ mutation with OptiPrime and tested these pegRNAs in mouse embryonic fibroblasts (MEFs) derived from *Kif1a*^lgdg^ mice by electroporating cells with *in vitro*-transcribed PE mRNA and chemically synthesized pegRNAs and nsgRNAs. While such a campaign would typically be cost-prohibitive due to the difficulty of synthesizing long RNA molecules, we hypothesized that the accuracy of OptiPrime would reduce the number of pegRNAs required to find an initial lead.

First, we evaluated the top eight silent edit combinations nominated by OptiPrime with only one RTT+PBS combination each (OP-1–8), which identified OP-5 as the pegRNA encoding the silent edit combination that yielded the highest editing efficiency of 10% (Figure 7B, 7C). In our second optimization assay, we evaluated seven additional RTT+PBS combinations with the same silent edit strategy as OP-5, revealing OP-5.4 as the lead pegRNA candidate that yielded 22% correction in heterozygous *Kif1a*^lgdg/+^ MEFs without any nsgRNA (Figure 7D). In total, these pegRNA optimization experiments required only 15 total pegRNAs—far fewer than the hundreds required for each edit in previous therapeutic PE studies^12,47^. With OP-5.4 in hand, we evaluated 15 nsgRNAs and identified several that enabled increases in editing efficiency exceeding two-fold. Although comprehensive evaluation of candidates in homozygous *Kif1a*^lgdg/lgdg^ MEFs was not possible due to low cell availability, we performed small-scale investigations in this system. We observed that some nsgRNA candidates induced high levels of indel formation in this setting, whereas nsgRNAs+65 and +114 maintained indel rates below 5%. We reasoned that these data, while limited, may more accurately reflect editing outcomes in the phenotypically relevant homozygous *Kif1a*^lgdg/lgdg^ model system. We therefore advanced both nsgRNAs +65 and +114, which in larger-scale characterization in heterozygous MEFs conferred 2.5- and 1.5-fold improvements in editing efficiency over PE2, yielding 39% and 23% editing, respectively.

Although nsgRNA +65 yielded higher editing efficiency, it also generated more indel byproducts compared to nsgRNA +114 (Figure 7E, Supplementary Figure 6A). Finally, we assayed both of these PE strategies with evolved PE6 prime editor variants^91^, which revealed that PE6b offers an additional 1.1-fold improvement over PEmax (Figure 7F), now achieving 64% (nsgRNA +65) and 51% (nsgRNA +114) editing efficiencies in heterozygous *Kif1a*^lgdg/+^ MEFs.

These results, obtained from only four experiments requiring a total of 4 weeks, demonstrate how OptiPrime can greatly accelerate the development of new, efficient therapeutic prime editing strategies. We packaged dual adeno-associated virus 9 (AAV9)^92^ with an epegRNA version of OP-5.4, either nsgRNA +65 or +114, and PE6b (Figure 7G). These AAV, along with an AAV encoding an EGFP–KASH transduction marker, were delivered *via* intracerebroventricular injection into heterozygous *Kif1a*^lgdg/+^ mice at birth (P0). After four weeks, mice treated with the AAV encoding both nsgRNAs showed above 40% average bulk brain cortex editing, with >70% editing efficiency among transduced (GFP^+^) cells (Figure 7H, Supplementary Figure 6B-C). Although we found lower editing levels in some brain regions, consistent with previous studies^47^, we found that over 50% of transcripts were edited in bulk tissue, suggesting that AAV9 transduction and prime editing occurs preferentially in cells that express Kif1a. Collectively, these findings show that OptiPrime can be used to rapidly generate therapeutic PE strategies to efficiently correct pathogenic mutations *in vivo*, including those that arise from *de novo* mutations that would often require *N*=1 treatments.

## Discussion

The versatility of PE offers immense potential as a tool for research and as a way to precisely tailor the genome of living systems for therapeutic and other applications. In this work, we demonstrated that OptiPrime, a mechanistic ML model, is able to predict PE efficiencies with high enough accuracy to greatly streamline an otherwise laborious pegRNA optimization process. Moreover, we showed that the mechanistic formulation underlying OptiPrime confers several advantages over traditional ML models, including the ability to use transfer learning and mechanism rewiring to predict outcomes of PE3 and twinPE, editing mechanisms not seen during training.

While OptiPrime was trained on many experiments in diverse cell types, we note that high-performing pegRNAs optimized using its HeLa cell parameters tend to also maintain their performance in HEK293T cells, while the converse is not true. Therefore, we recommend that researchers interested in designing pegRNAs with OptiPrime use its default settings for typical applications. This recommendation is supported by our encouraging *in vitro* results in primary human T cells, patient-derived fibroblasts, and mouse embryonic fibroblasts, as well as *in vivo* mouse brain editing, all contexts in which the pegRNAs generated using OptiPrime’s default parameters performed well.

OptiPrime performs less accurately in cases in which Cas9 nuclease scores fail to correlate with PE efficiency (Figure 4G). Similarly, during our exploratory data analysis of PE3 data, we showed that Cas9 nuclease scores fail to predict nsgRNA performance (Figure 5C). In future studies, we hope to further improve the accuracy of OptiPrime by replacing Rule Set 3 scores with PE-specific heuristics for protospacer efficiency. Moreover, we speculate that further studies elucidating the DNA repair mechanisms governing PE3, PE3b, and twinPE can lead to improved performance when selecting optimal nsgRNAs and twinPE pegRNA pairs.

Similar to other models that predict PE efficiency based on pegRNA sequences, OptiPrime was primarily trained on data from high-throughput PE experiments performed on a library of synthetic reporters randomly integrated into the genome. Information about chromatin context is therefore not incorporated into OptiPrime. Researchers interested in augmenting OptiPrime predictions with chromatin state information can use models such as ePRIDICT, which predicts relative editing efficiency at genomic loci in K562 cells without considering pegRNA sequences. Future studies may integrate chromatin state information into OptiPrime directly, although the cell type dependence of chromatin state may limit the generality of such an approach.

We found that empirical editing efficiencies cluster based on pegRNA similarity. While a deep exploration of this phenomenon to further increase the efficiency of navigating OptiPrime-nominated pegRNA space is the subject of ongoing efforts, and comparisons in this work were made simply by taking the top ranked outputs from OptiPrime, we expect that future work may integrate OptiPrime with iterative pegRNA optimization using techniques such as Bayesian optimization to further streamline the development of PE strategies by minimizing the number of pegRNAs tested to achieve efficient prime editing.

## Data availability

Raw high-throughput sequencing data have been deposited at the NCBI Sequence Read Archive (SRA) under accession PRJNA1314411 and are publicly available as of the date of publication. This paper analyzes existing, publicly available high-throughput sequencing data, accessible on the SRA (PRJNA735408, PRJNA1055086, PRJNA1211588). Plasmids generated in this study will be available from Addgene as of the date of publication. Requests for other resources should be directed to David R. Liu.

## Code availability

All original code relevant to training and modeling will be publicly available as of the date of publication under a non-commercial license. Any additional information required to reanalyze the data reported in this paper is available from the lead contact upon request.

## Acknowledgements

Funding was provided by US NIH grants U01AI142756 (D.R.L.), RM1HG009490 (D.R.L), R01EB022376 (D.R.L.), R35GM118062 (D.R.L.), R01HL147324 (M.J.O.), P01CA065493 (M.J.O.), R01AR063070 (M.J.O.), DP2CA281401 (B.P.K.), and P01HL142494 (B.P.K.), the Howard Hughes Medical Institute, the Bill & Melinda Gates Foundation. This work was supported by the NIH IGNITE Program (R61NS133266-02 to M.T. and C.M.L.). The authors acknowledge funding from NSF Graduate Research Fellowships to A.H., P.J.C., A.C., A.A.S., and M.W.S., and a Natural Sciences and Engineering Research Council of Canada (NSERC) Postgraduate Scholarship-Doctoral (PGS D – 567791) to R.A.S. The authors thank Peyton Randolph, Jordan Doman, and Żaneta Matuszek for helpful discussions and John Harvey and Benjamin Deverman for technical assistance. We thank the Jackson Laboratory (JAX), all RDTC (Rare Disease Translational Center) and JCPG (JAX Center for Precision Genetics) members for their operational support.

## Author information

A.H., P.J.C., and D.R.L conceptualized the project. A.H., P.J.C., A.H.L., A.N.M., B.J.S., M.J.O., C.F.H., M.T., C.N., V.S., A.C., P.Z.C., X.D.G., A.A.S., H.A.S., and R.A.S. performed experiments and analyzed the data. B.R.B., J.T., M.J.O., C.M.L., and D.R.L. supervised the project. A.H., I.M., and R.K.K wrote the software. A.H., P.J.C., A.H.L., M.T., C.M.L., and D.R.L. wrote the manuscript with input from all authors. A.H., P.J.C., and D.R.L. have filed patent applications on aspects of this work. A.H. is currently an employee at Chan Zuckerberg Biohub. P.J.C. is currently an employee at Prime Medicine. C.F.H. is currently an employee at the Cystic Fibrosis Foundation. X.D.G. currently holds an academic appointment at the University of Pittsburgh. S.P. is currently an employee at CRISPR Therapeutics. R.K.K. is currently an employee at Google Deepmind. M.W.S. is currently an employee at Genentech. D.R.L. is a consultant and/or equity holder of Prime Medicine, Beam Therapeutics, Pairwise Plants, and nChroma Bio, companies that use and/or deliver gene editing or genome engineering technologies. R.A.S. and B.P.K. are inventors on patent applications filed by Mass General Brigham (MGB) that describe genome engineering technologies. B.P.K. is a consultant for EcoR1 capital, Novartis Venture Fund, and Jumble Therapeutics, and is on the scientific advisory boards of Acrigen Biosciences, Life Edit Therapeutics, and Prime Medicine. B.P.K. has a financial interest in Prime Medicine, Inc., a company developing therapeutic CRISPR-Cas technologies for gene editing. B.P.K.’s interests were reviewed and are managed by MGH and MGB in accordance with their conflict-of-interest policies. M.J.O. receives compensation as a consultant for Agathos Biologics. B.R.B. reports research funding from BlueRock Therapeutics and Carisma Therapeutics and consulting fees from BlueRock Therapeutics, Editas Medicine, Janssen Oncology, Sandoz, Legend Biotech, GentiBio Inc., and Magenta Therapeutics. The remaining authors declare no conflicts of interest.

## Methods

### Culture conditions for immortalized cell lines

HEK293T and HeLa cells were cultured in Dulbecco’s Modified Eagle Medium (DMEM) plus GlutaMAX (Thermo Fisher) supplemented with 10% fetal bovine serum (FBS) (Thermo Fisher). Both cell types were passaged every 2–3 days, maintained below 80% confluency, cultured at 37 °C with 5% CO_2_, and tested negative for mycoplasma.

### Isolation of and culture conditions for primary human fibroblasts

With IRB approval, informed patient consent, and in accordance the Declaration of Helsinki Principles, a punch biopsy was obtained from a patient with recessive dystrophic epidermolysis bullosa (RDEB). The tissue was minced and primary fibroblasts were expanded and subsequently propagated in Alpha MEM and 1× final concentrations of non-essential amino acids, penicillin, streptomycin, GlutaMAX, and anti-oxidant supplement (all from Thermo Fisher Scientific), 20% fetal bovine serum (Atlas Biologicals, Fort Collins, CO), 0.5 ng/mL epidermal growth factor, and 10 ng/mL epidermal growth factor (each from Sigma-Aldrich, St Louis, MO).

### Isolation of and culture conditions for primary human T cells

Blood units were obtained from unrelated donors (*n* = 3) without identifying information from Memorial Blood Centers (Saint Paul, MN). Peripheral blood mononuclear cells (PBMCs) were isolated from buffy coats using Lymphoprep (STEMCELL Technologies) and SepMate tubes (STEMCELL Technologies) and cryopreserved in CryoStor CS10 Cell Cryopreservation Media (STEMCELL Technologies). T-cells were isolated from PBMCs using the EasySep Human T Cell Isolation Kit (STEMCELL Technologies) and activated with Dynabeads Human T-Expander CD3/CD28 at a 3:1 bead to cell ratio in complete T-cell media comprised of: X-VIVO 15 (Lonza), 10% Human Ab serum (Valley Biomedical), 1× GlutaMAX, 1× penicillin/streptomycin (Life Technologies), 1× *N*-acetyl-L-cysteine (Sigma-Aldrich) and 300 IU/mL interleukin-2 (PeproTech).

### Isolation of and culture conditions for mouse embryonic fibroblasts

To obtain mouse embryonic fibroblasts (MEFs), mice embryos were collected from timed matings of heterozygous JR #16894 C3.Cg-*Kif1a*^lgdg^/GrsrJ x JR #664 C57BL/6J between embryonic day (E) E12.5 and E14.5. Embryos were dissected out of uterine horns, washed in 70% ethanol, and placed in a 10 cm petri dish with PBS. Embryos were separated from the placenta, head, liver, and heart and blood clots were washed out. Embryo cells were minced with a razor blade and dissociated with 2 mL of trypsin-EDTA and incubated at 37 °C for 10 minutes with shaking. Cells were resuspended with an additional 4 mL of DMEM (Thermo Fisher) supplemented with 10% embryonic stem cell FBS (Thermo Fisher) and vigorously resuspended. Any residual debris was removed by centrifugation and cells were plated in a T75 flask. MEFs were maintained in DMEM with GlutaMAX (Thermo Fisher), PSN (Thermo Fisher), and 10% embryonic stem cell fetal bovine serum (Thermo Fisher) at 37 °C in 5% CO2. Media was changed after 48 h. Four days post-isolation, cells were washed with PBS, trypsinized (0.05% Trypsin-EDTA), and frozen at passage P1 in cell culture media supplemented with 10% DMSO, 5.0×10^6^ cells/mL per cryo-vial. Cryo-preserved P1 MEFs were thawed and used for *in vitro* gene editing experiments.

### Paired pegRNA–target site library cloning

A lentiviral transfer plasmid for library cloning (pPC1535) was designed to contain a human U6 promoter–2×BsmBI and a PuroR–T2A–BFP marker under expression from an EF1α promoter (pEF1α). An oligonucleotide library of paired pegRNAs (containing a 2×BsmBI sequence instead of the sgRNA scaffold) and target sites (Twist Biosciences) was PCR amplified with Kapa HiFi Master Mix DNA Polymerase (Roche) and purified with the QIAquick PCR Purification Kit (QIAGEN). Backbone plasmid (pPC1535) was digested with BsmBI-v2 (New England Biolabs), treated with Quick CIP (New England Biolabs), then gel purified on a 1% agarose gel and purified with 0.75× SPRIselect beads (Beckman Coulter). Digested pPC1535 and amplified oligonucleotide libraries were then cloned with Gibson assembly with NEBuilder HiFi DNA Assembly (New England Biolabs) and purified with 1× SPRIselect beads (Beckman Coulter). Next, assembled plasmids were electroporated into 10-beta electrocompetent *E. coli* (New England Biolabs) and plated on LB agar plates with 50 µg/mL carbenicillin. After cells were grown for ∼16 h at 37 °C, colonies were scraped and plasmid DNA was isolated with the Plasmid Plus Midi Kit with endotoxin removal (QIAGEN).

For the second cloning step, in which the pegRNA scaffold was incorporated into the U6-driven pegRNAs, midiprepped plasmid was digested with BsmBI-v2 (New England Biolabs) and gel purified on a 1% agarose gel. Next, phosphorylated oligos encoding the BlpI-modified SpCas9 F+E sgRNA scaffold were ligated to digested plasmid with T4 DNA ligase (New England Biolabs) in the presence of Esp3I (ThermoFisher), then purified with 1× SPRIselect (Beckman Coulter). Assembled plasmids were electroporated into 10-beta electrocompetent *E. coli* (New England Biolabs) and plated on LB agar plates with 50 µg/mL carbenicillin. After cells were grown for ∼16 h at 37 °C, colonies were scraped and plasmid DNA was isolated with the Plasmid Plus Midi Kit with endotoxin removal (QIAGEN).

### Lentivirus production, functional titering, and transduction

The day before transfection, 10×10^6^ HEK293T cells were seeded in 15-cm dishes with DMEM (ThermoFisher) supplemented with 10% FBS (Millipore Sigma). When cells reached 60% confluency (∼16 h after seeding), cells were transfected with 13.3 µg lentiviral transfer plasmid, 6.7 µg pMD2.G (Addgene #12259), and 10 µg psPAX2 (Addgene #12260) with 120 µL Lipofectamine 2000 (ThermoFisher) according to the manufacturer’s protocol.

6 h post transfection, media was exchanged with fresh DMEM (ThermoFisher) supplemented with 10% FBS (Millipore Sigma). 48 h after transfection, viral supernatant was centrifuged at 3000× g for 15 min to remove cellular debris, filtered through a 0.45 mm PVDF filter (Corning), aliquoted, and stored at −80 °C. 2×10^6^ HEK293T or HeLa cells were “spinfected” with 0, 10, 20, 50, 100, or 200 µL of a thawed aliquot of virus in the presence of 10 µg/mL polybrene (Sigma Aldrich) in 6-well plates at 1000× g for 120 min, then transferred to an incubator at 37 °C with 5% CO_2_. 24 h post-infection, cells were trypsinized, resuspended in PBS, and fixed with 4% w/v paraformaldehyde (Sigma Aldrich). Fixed cells were strained into a round bottom test tube with a 35 mm strainer cap (Falcon) and BFP expression was quantified with a CytoFlex LX cytometer to calculate the MOI.

To make the final cell libraries, cells were transduced with a volume of virus that would result in ∼0.2 MOI with 1,000× coverage of each library member post-selection. 1 day after infection, cells were treated with 1 µg/mL puromycin (ThermoFisher) to select for cells with integrated library members 2 days after infection, an additional 2 µg/mL puromycin was added to cells.

### Paired pegRNA–target site library screens

All screens were performed in biological duplicate and with PE2 and PE4 conditions on cells that had undergone at least 6 days total of puromycin selection. A single replicate of untreated cells was always sequenced for background subtraction of editing efficiency. The day before transfection, 10×10^6^ HEK293T cells or 3×10^6^ HeLa cells were seeded in DMEM (ThermoFisher) supplemented with 10% FBS (Millipore Sigma) into 15 cm dishes. Cells were transfected at 70% confluency (∼16 h after seeding).

Each 15-cm dish of HEK293T cells was transfected with 6.7 µg of pCMV–PEmax and 3.3 µg of either pEF1α–RFP (PE2) or pEF1α–hMLH1dn (PE4) with 100 µL of Lipofectamine 2000 (ThermoFisher) according to manufacturer’s protocols. Each 15 cm dish of HeLa cells was transfected with 25 µg of pCMV–PEmax–P2A–BSD and 25 µg of either pEF1α–RFP (PE2) or pEF1α–hMLH1dn (PE4) with 140 µL of TransIT-HeLa reagent (Mirus Bio; no HeLaMONSTER reagent used) according to manufacturer’s protocols. 24 h following transfection of HeLa cells, cells were treated with 10 µg/mL blasticidin S (ThermoFisher) to select for expression of PEmax. 48 h following transfection of HeLa cells, the media was replaced with DMEM (ThermoFisher) supplemented with 10% FBS (Millipore Sigma), 1× Penicillin/Streptomycin (ThermoFisher), 10 µg/mL blasticidin S (ThermoFisher), and 1 µg/mL puromycin. 72 h following transfection of HEK293T cells or 120 h following transfection of HeLa cells, cells were trypsinized, pelleted, and stored at −80 °C.

### High-throughput sequencing of pooled screens

Genomic DNA was extracted from all screen cells with the NucleoSpin Blood XL Maxi kit (Machery-Nagel). The entirety of the genomic DNA from each screen condition was used in the initial round of PCR (PCR1) to amplify the region containing the gRNA and target site.

Each 100 mL PCR1 was performed with 5 µg of genomic DNA as template, 1 mM of each of Lib-FWD and Lib-REV, and 50 mL of NEBNext Ultra II Q5 Master Mix (New England Biolabs) on a BioRad C1000 thermal cycler with the following cycling parameters:

98 °C for 30 s,

22 cycles of

98 °C for 10 s,

65 °C for 75 s,

65 °C for 5 min.

The reactions were characterized by Agilent TapeStation. 1 mL of pooled PCRs1 from each condition were gel purified (QIAGEN) and further purified with 0.8× SPRISelect. A following PCR step (PCR2) enabled indexing of the samples through addition of barcodes and binding to the flow cell through addition of the P5/P7 Illumina sequences.

For each screen condition, 4×50 µL PCRs2 were performed. Each PCR2 was performed with 10 ng of purified PCR1 amplicon was used as template, 600 nM each of a PE-FWD primer and a PE-REV primer, and 25 µL of Kapa HiFi Master Mix DNA Polymerase (Roche) on a BioRad C1000 thermal cycler with the following cycling parameters:

98 °C for 3 min,

8 cycles of

98 °C for 20 s,

65 °C for 15 s,

72 °C for 15 s,

72 °C for 1 min.

The reactions were characterized by Agilent TapeStation. Reactions were combined and purified with 0.8× SPRISelect, then normalized for concentration prior to pooling and sequencing. Libraries were sequenced on an Illumina NovaSeq 6000 with the S1 Reagent Kit v1.5 with 50 cycles for R1, 8 cycles for I1, 8 cycles for I2, and 150 cycles for R2.

### Processing of pegRNA–target site library screen sequencing data

Sequencing data from the NovaSeq 6000 were demultiplexed into fastq.gz files for each individual screen based on the index reads with bcl2fastq2. In these data, R1 reads cover the pegRNA spacer, while R2 reads contain the sequence outcome resulting from PE as well as a barcode specifying pegRNA identity. Within each screen, R2 reads were demultiplexed into individual fastq.gz files based on the barcode sequence in R2. If the sequenced barcode did not match the sequenced spacer (from R1), then the read was discarded. These individual R2 fastq.gz files were then analyzed individually with CRISPResso2 ^93^ to obtain editing and indel frequencies for each pegRNA. For CRISPResso analysis, R2 read sequences were aligned to a reference sequence in HDR mode with a quality cutoff of Q30. For each amplicon, the CRISPResso2 quantification window was positioned to include the entire sequence between the nick site and homology end with 10 additional bp of buffer on either side. Frequencies for these edits were quantified as: (number of HDR-aligned reads) / (number of reads aligned to any amplicon). Frequencies for indels were quantified as: (number of indel-containing reads) / (number of reads aligned to any amplicon).

### Preparation of mRNA

Prime editor mRNA was generated using *in vitro* transcription (IVT). Briefly, the prime editor transcript, containing a 5′ untranslated region (UTR), Kozak sequence, prime editor open reading frame, and 3′ UTR were PCR amplified from a template plasmid containing an inactive T7 (dT7) promoter. PCR primers repaired this dT7 promoter and also installed a 119 nt poly(A) tail. The purified PCR double-stranded DNA amplicon was used as an IVT template using the HiScribe T7 high-yield RNA synthesis kit (NEB). IVT was performed following the manufacturer’s optional protocol to include CleanCap reagent AG (Trilink) and substitute N^1^-methylpseudouridine-5′-triphosphate (Trilink) for uridine triphosphate, after which IVT template was degraded with DNase I (NEB). A Monarch Spin RNA Cleanup Kit (NEB) was used to purify mRNA from complete IVT reactions, and mRNA transcripts were reconstituted in nuclease-free water.

### Preparation of AAV

Transfer vectors were cloned in a v1em twin prime editor AAV architecture^92^ adapted for the PE6b editor variant. Briefly, synthetic gene fragments (IDT) with appropriate restriction recognition sites and containing an nsgRNA cassette driven by a mouse U6 promoter and an epegRNA cassette driven by a human U6 promoter were installed into the C-terminal transfer plasmid via restriction digestion and T4 ligation (NEB). Plasmids were transformed into chemically competent NEB Stable cells (NEB), which were plated on LB agar plates with 50 µg/mL carbenicillin and grown at 30 °C overnight. Single colonies were picked and grown in 2xYT media with 50 µg/mL carbenicillin for 20 hours, and plasmids were isolated using a ZymoPURE II Plasmid Maxiprep Kit (Zymo Research).

Recombinant AAVs (AAV9) were produced in suspension HEK293T cells, using F17 media (ThermoFisher). Cell suspensions were incubated at 37 °C, 8% CO2, 80 RPM. 24 hours before transfection, cells were seeded in 500–1000 mL of media at ∼1×10^6^ cells/mL. The day after, cells (∼2×10^6^ cells/mL) were transfected with pHelper, pRepCap and pTransgene (2:1:1 ratio, 2 µg DNA per million cells) using Transporter 5 transfection reagent (Polysciences) with a 2:1 reagent:DNA ratio. Three days post-transfection, cells were pelleted via centrifugation at 2000 RPM for 12 minutes in Nalgene conical bottles. The supernatant was discarded, and cell pellets were stored at −20 °C until purification. Each pellet, corresponding to 500 mL of cell culture, was resuspended in 14 mL of 500 mM NaCl, 40 mM Tris-base, 10 mM MgCl2, with Salt Active Nuclease (ArcticZymes) at 100 U/mL. Afterwards, the lysate was clarified at 5000 RCF for 20 minutes and loaded onto a density step gradient containing OptiPrep (Cosmo Bio) at 60%, 40%, 25%, and 15% at a volume of 6, 6, 8, and 5 mL respectively in OptiSeal tubes (Beckman Coulter). The step gradients were spun in a Beckman Type 70ti rotor (Beckman Coulter) in a Sorvall WX+ ultracentrifuge (ThermoFisher) at 67,000 RPM for 1 hour and 15 minutes at 18 °C. Afterwards, ∼4.5 mL of the 40–60% interface was extracted using a 16-gauge needle, filtered through a 0.22 μm PES filter, buffer exchanged with 100K MWCO protein concentrators (ThermoFisher) into PBS containing 0.001% Pluronic F-68, and concentrated down to a volume of 200–1000 μL. The concentrated virus was filtered through a 0.22 μm PES filter and stored at 4 °C.

Titers of purified AAVs were measured using an AAVpro Titration Kit (for Real Time PCR) Ver. 2 (Takara) with 1:50,000 final dilutions of AAV stocks as input, following the manufacturer’s protocol. Viral stocks were concentrated to a minimum of 2×10^13^ vg/mL using Pierce PES 100K MWCO Concentrators (ThermoFisher) pre-equilibrated with 0.001% F-68 in PBS (ThermoFisher) and sterile filtered using 0.22 µm GV Durapore centrifugal filter units (Millipore). Concentrated viral titers were confirmed using an AAVpro Titration Kit (for Real Time PCR) Ver.2 (Takara) with 1:50,000 final dilutions of AAV stocks as input, following the manufacturer’s protocol. Samples were sent to Plasmidsaurus for verification of AAV genome integrity via Oxford Nanopore long-read sequencing.

### Transfection and genomic DNA processing of HEK293T cells

The day before transfection, approximately 16,000 HEK293T cells were plated into each well of a 96-well plate (Corning) in DMEM (ThermoFisher) supplemented with 10% FBS (Millipore Sigma). When cells reached 70% confluency (∼18 h after plating), cells were transfected with variable amounts of plasmid DNA and 0.5 µL of Lipofectamine 2000 (ThermoFisher) diluted in Opti-MEM I (ThermoFisher) according to manufacturer’s protocols. For a typical 96-well transfection using standard prime editor systems, the following plasmid amounts were transfected:

- PE2: 200 ng prime editor and 50 ng pegRNA
- PE3: 200 ng prime editor, 50 ng pegRNA, and 15 ng nsgRNA
- PE4: 200 ng prime editor, 100 ng MLH1dn, and 50 ng pegRNA
- PE5: 200 ng prime editor, 100 ng MLH1dn, 50 ng pegRNA, and 15 ng nsgRNA
- eePASSIGE: 100 ng prime editor, 100 ng Bxb1, 150 ng donor, 15 ng pegRNA 1, 15 ng pegRNA 2

Following transfection, cells were incubated at 37 °C with 5% CO2 for 72 h before lysis and genomic DNA harvest.

Genomic DNA was isolated from HEK293T cells following a custom lysis protocol and paramagnetic bead extraction. To lyse cells, growth medium was carefully removed from cell culture plates and 100 µL of lysis buffer (100 mM Tris-HCl pH 8.0, 200 mM NaCl, 5 mM EDTA, 0.05% SDS, 4.0 mg/mL proteinase K (QIAGEN), and 12.5 mM dithiothreitol) was added to each well of a 96-well plate. Plates with lysis buffer were incubated at 55 °C for 12–20 h with shaking. To extract genomic DNA, one volume of lysate was thoroughly mixed with one volume of Ampure XP paramagnetic beads (Beckman Coulter) and incubated for 5 min, separated on plate magnet, washed with 70% ethanol three times with bead resuspension, and eluted in 50 µL of nuclease-free water.

### Electroporation of primary human fibroblasts

On the day of electroporation, the fibroblasts were trypsinized (final concentration of 0.05% in 1 mM EDTA/PBS) and enumerated using a Countess Cell Counter (Thermo Fisher Scientific). 100,000 cells were mixed with 1 µg of PEmax-encoding mRNA, 250 ng GFP-encoding mRNA, and 90 pmol of pegRNA. Cells were electroporated with the Neon NxT Electroporation System (Thermo Fisher Scientific) using a 10 µL tip using one 20 ms pulse of 1700 V. Cells then were plated in a 24 well plate in complete media and harvested four days later for genomic DNA isolation with the Monarch Spin gDNA Extraction Kit (New England Biolabs).

### Electroporation of primary human T cells

48h post stimulation, T cell activation beads were removed and the cells were replated for ∼2 hours in T-cell media. Cells were harvested, counted, and 300,000 T cells were mixed with 1 µg of PEmax-encoding mRNA and 90 pmol of pegRNA. Cells were electroporated with the Neon NxT Electroporation System (Thermo Fisher Scientific) with 10 µL tips and Buffer T and E10 using three 10 ms pulses of 1400 V. Cells were then plated in a 48 well plate with 600 µL of complete T-cell media and then harvested at 72 hours by centrifugation and resuspended in QuickExtract DNA Extraction Solution (Biosearch Technologies). Plates with lysis buffer were incubated at 65 °C for 10 m with shaking.

### Electroporation of mouse embryonic fibroblasts

Mouse embryonic fibroblasts were trypsinized at 70% confluency, counted, and washed with PBS. Per electroporation, 200,000 cells were resuspended into 20 µL of SE nucleofector solution with SE supplement solution (Lonza) containing 1 µg of prime editor mRNA, 90 pmol of synthetic pegRNA (IDT), and 60 pmol synthetic nsgRNA (Synthego) (if applicable). Cells were then electroporated with a 4D-Nucleofector with X Unit (Lonza) using program CM-130 and the SE Cell Line 4D-Nucleofector X Kit S (Lonza). Cells were allowed to recover with addition of 80 µL of prewarmed DMEM for 10 minutes then plated in a 24-well plate with 500 µL DMEM and collected after 72 hours by addition of lysis buffer.

### Mouse strains

The leg dragger mutation (Kif1a L181F) arose as a spontaneous mutation on a subline of C3.NB-H2 mice (The Jackson Laboratory, JR #16894, MGI:5752896) and referred to as C3.Cg-*Kif1a*^lgdg^/GrsrJ mice. The lgdg mutation is maintained on a hybrid genetic background with C57BL/6J (The Jackson Laboratory, JR #664, B6J;C3-Kif1a^lgdg^/Lutzy). For the generation of B6J;C3-*Kif1a*^lgdg^ mice, heterozygous C3.Cg-*Kif1a*^lgdg^/GrsrJ mice (JR #16894) were intercrossed with C57BL/6J mice (JR #664). Mice were genotyped at birth, wean age (∼postnatal P28), and triple confirmed when reaching study end (e.g., necropsy for tissue collection). The genotyping primers are listed below and are applicable regardless of the genetic background. Furthermore, B6J;C3-Kif1a^lgdg^ embryos (small piece of the tail, day of MEF isolation) and MEFs (small aliquot of cells during passaging and cryo-preservation) were genotyped using the same genotyping assay. The Jackson Laboratory Animal Care and Use Committee approved all mouse protocols.

For animal experiments, simplified genotype abbreviations, such as *Kif1a^+/+^* (B6J;C3-*Kif1a*^+/+^) and *Kif1a*^lgdg/+^ (B6J;C3-*Kif1a*^lgdg/+^) mice, were used throughout.

Genotyping Primers (lgdg mutation):

- Common Forward: 5′-ATGGAGATTTACTGTGAGCGAG-3′
- Common Reverse: 5′-GGTCCTGGATGTCATTGTAGG-3′
- WT Probe (HEX): 5′-TGTGGAGGACCTCTCCAAGCT-3′
- lgdg mutant Probe (FAM): 5′-TGTGGAGGACTTCTCCAAGCT-3′

### Intracerebroventricular injection

For neonatal ICV injection (targeting of brain ventricles located below the cortex), heterozygous C3.Cg-*Kif1a*^lgdg^/GrsrJ (JR #16894) males were crossed with C57BL/6J (JR #664) females. C57BL/6J females were singly housed 10-12 days post mating set up. The week of birth, females were checked daily for the birth of the litter. On the day of birth (postnatal day P0), the litter of B6J;C3-*Kif1a*^lgdg^ mice was injected, and operators were blinded to genotype and the treatment (test articles AAV9-OP5.4-+65 and AAV9-OP5.4-+114). For gene editing studies, mice were bilaterally injected with either prime editor (total dose: 1.1×10^11^ vg/mouse, consisting of 5×10^10^ vg each N- and C-terminal vector and 1×10^10^ vg EGFP-KASH vector, 2.5µl per hemisphere). Uninjected B6J;C3-*Kif1a*^lgdg^ mice were used as a control group for each prime editor. Post-injection, B6J;C3-*Kif1a*^lgdg^ pups were monitored daily for survival.

### Necropsy and tissue collection

Mice were euthanized via CO_2_ asphyxiation 28 days after birth. Brain tissues were harvested and dissected into cortex, brainstem, hippocampus, and cerebellum. Samples were frozen on dry ice and stored at −80 °C until further analysis.

### Nuclear isolation, sorting, nucleic acid extraction, and cDNA synthesis

Frozen brain tissues were resuspended in 2 mL of Nuclei Extraction Buffer (Miltenyi Biotec) supplemented with 0.2 U/µL Murine RNase Inhibitor (NEB) and transferred to gentleMACS C Tubes (Miltenyi Biotec). Samples were homogenized using the default program “4C_nuclei_1” on a gentleMACS Octo Dissociator (Miltenyi Biotec). The nuclei suspension was washed with an additional 2 mL of Nuclei Extraction Buffer, filtered through a 100 µm MACS SmartStrainer (Miltenyi Biotec) into a 5 mL centrifuge tube, and centrifuged at 500×g at 4 °C for 5 minutes. The supernatant was aspirated, and the pelleted nuclei were resuspended in 4 mL of ice-cold nuclei suspension buffer consisting of PBS supplemented with 100 µg/µL albumin (NEB), 3.33 µM Vybrant DyeCycle Ruby Stain (Thermo Fisher), and 0.2 U/µL RNase Inhibitor, Murine (NEB). Samples were centrifuged again at 500×g at 4 °C for 5 minutes, and the supernatant was aspirated. The pellet was resuspended in 1 mL of ice-cold nuclei suspension buffer and filtered through a 35 µL cell strainer into a 5 mL Falcon tube (Corning). Nuclei were sorted using a Sony SH800 Cell Sorter (Sony Biotechnology) with a 100 µm chip in the Broad Institute Technology Space. Singlet DyeCycle Ruby-positive events were collected and back-gated to establish forward and side scatter area gates for nuclei. Nuclei were then sorted based on GFP fluorescence to yield “bulk” (containing all nuclei irrespective of GFP expression) and “GFP-positive” populations. For each population, 25,000 nuclei per tissue sample were sorted into 540 µL of Buffer RLT Plus from the AllPrep DNA/RNA Mini kit (QIAGEN) supplemented with 74 mM DTT. DNA and RNA were purified according to the manufacturer’s protocol. The SuperScript IV First-Strand Synthesis System with ezDNase was used with kit-supplied random hexamers to generate cDNA from 9 µL of RNA input, following the manufacturer’s protocol. Synthesis reactions were cleaned up using SPRIselect (Beckman Coulter) at a 1.8x bead ratio and eluted in 35 µL of nuclease-free water. For sequencing analysis, 10 µL of purified cDNA was used as input for PCR1.

### High-throughput sequencing of genomic loci

To assess gene editing, loci were amplified from genomic DNA samples with two rounds of PCR and deep sequenced. Briefly, an initial PCR step (PCR1) amplified the genomic sequence of interest with synthetic primers (IDT) containing Illumina forward and reverse I5/I7 adapters. Each 20 µL PCR1 reaction was performed with 500 nM of each primer, 0.8 to 1.0 µL of genomic DNA, 1× SYBR Green I, and 10 µL Q5 High-Fidelity 2× Master Mix (New England Biolabs) on a CFX96 Touch Real-Time PCR detection System with the following thermocycling conditions:

98 °C for 3 min,

30 cycles of

98 °C for 20 s,

65 °C for 15 s,

72 °C for 15 s,

72 °C for 2 min

### PCR1 reactions were monitored with SYBR Green I fluorescence to avoid over-amplification

The subsequent PCR step (PCR2) added unique i7 and i5 Illumina barcode combinations to both ends of the PCR1 DNA fragment to enable demultiplexing. Each PCR2 was performed with 10 ng of purified PCR1 amplicon was used as template, 600 nM each of a PE-FWD primer and a PE-REV primer, and 25 µL of Kapa HiFi Master Mix DNA Polymerase (Roche) on a BioRad C1000 thermal cycler with the following cycling parameters:

98 °C for 3 min,

8 cycles of

98 °C for 20 s,

65 °C for 15 s,

72 °C for 15 s,

72 °C for 1 min

PCR2 products were pooled by common amplicons, separated by electrophoresis on a 1% agarose gel, purified using the QIAquick Gel Extraction Kit (QIAGEN), and eluted in nuclease-free water. DNA amplicon libraries were quantified with a Qubit 3.0 Fluorimeter (ThermoFisher), then normalized, pooled, and sequenced using the Illumina MiSeq Reagent Kit v2 with 280–300 single-read cycles.

### Quantification of editing in amplicon sequencing data

Sequencing reads were demultiplexed with bcl2fastq2. Amplicon sequences were aligned to a reference and edited sequence with CRISPResso2^93^ in HDR mode using the parameters -q 30 --discard_indel_reads. For each amplicon, the CRISPResso2 quantification window was positioned to include the entire sequence between pegRNA- and (where applicable) sgRNA-directed Cas9 cut sites as well as an additional 10 bp beyond both cut sites. All prime editing efficiencies describe percentage of (number of HDR-aligned reads) / (number of reads aligned to any amplicon).

All prime editing indel rates describe percentage of (number of aligned reads discarded) / (number of reads aligning to any amplicon).

### Design of paired pegRNA–target site libraries

For both Lib-MMR and Lib-CV, we designed pegRNAs that we hypothesized would give high editing efficiencies while matching the criteria in Tables 1 and 2. For pegRNAs that encode substitutions, we used an initial RTT homology length of 9 + *L*, where *L* denotes the number of substituted bases. For pegRNAs that encode insertions or deletions, we used an initial RTT homology length of 19 + *L*, where *L* is the absolute length difference between the unedited and edited sequences. These initial RTT lengths were then altered (if necessary) to find the closest RTT length such that the first base of the RTT would not be a cytosine. All pegRNAs used PBS length 13, since we hypothesized that this PBS length would provide sufficient signal to learn MMR preferences while not occupying excessive library space.

### Model implementation and training

OptiPrime and all related models were implemented in Python 3.11 with jax (v.0.4.30) and flax (v.0.10.2). Stochastic gradient descent was performed with optax (v.0.2.4) using the AdamW optimizer with a linear warm-up ramp over 25 epochs to an initial learning rate of 0.001, then exponentially decaying by a factor of 0.99 each epoch for 75 epochs. Model hyperparameters were optimized manually with 5-fold stratified cross-validation. For detailed modeling considerations, readers are encouraged to read Supplemental Text 1.

We used code provided by each respective initial report to evaluate retrospective predictions made by PRIDICT2.0 and DeepPrime. In each case, we chose the model version most appropriate for the cellular system being studied, as listed in the caption for the respective figures. We validated that our code produced the correct outputs by comparing to outputs from each model’s respective webserver.

PE3 models were implemented in Python 3.11 with xgboost (v.3.0.0). In addition to outputs from OptiPrime, the following hand-crafted PE3-specific features were implemented and used:

- nick_offset: Distance (bp) between pegRNA and nsgRNA nicks
- hom_offset: Distance (bp) between end of RTT homology arm and nsgRNA nick
- PE3b: Whether the nsgRNA spacer only matches its target after flap synthesis
- pe_type: Whether PE3 or PE5 is used
- cell_type: Categorical encoding of the cell type used

### Selection of pegRNAs for prospective use cases

To best simulate prospective use cases, we entered the mutation that we wanted to correct into OptiPrime, PRIDICT2.0, or DeepPrime. For all experiments with OptiPrime, we used our default settings (HeLa cells; PE2max; epegRNAs) and took the top pegRNAs as ordered by silent edit combination. For experiments with PRIDICT2.0, we selected outputs as ranked by HEK score for experiments performed in HEK293T cells and as ranked by K562 score for all other experiments. For experiments with DeepPrime, we selected outputs as ranked by HEK293T_PE2max-e for experiments performed in HEK293T cells and as ranked by A549_PE2max-e for all other experiments, since the original authors only performed screens in A549 cells with both PE2max and epegRNAs.

### Quantification and statistical analysis

Correlation coefficients were calculated in Python with the scipy.stats package (v.1.14.1). Unless otherwise noted, *r* denotes a Pearson (linear) correlation and ρ denotes a Spearman rank correlation. Accordingly, we compute and report *r* when assessing quantitative predictive accuracy between correlates and ρ when assessing the ability to rank-order outcomes. *P*-values reported for both types of correlation coefficient represent the two-tailed significance of observing a correlation of the given magnitude under Student’s *t*-distribution with *n* − 2 degrees of freedom. We used the two-sided Steiger’s *Z*-test^94^ to compute *P*-values of differences between correlations on identical datasets to compare model performances. The number of independent biological replicates and technical replicates for each experiment are described in either the figure legends or the Methods section. Unless otherwise specified, all error bars represent standard deviation.

