## Supplemental Information for "Mechanistic machine learning enables interpretable and generalizable prediction of prime editing outcomes"

**Supplementary Text 1.** OptiPrime design considerations.

**Supplementary Figure 1.** A mechanism of prime editing in mammalian cells.
**Supplementary Figure 2.** High-throughput lentiviral screens recapitulate the sequence determinants of mammalian mismatch repair (MMR)
**Supplementary Figure 3.** The HetFormer is a neural network designed to model DNA heteroduplexes.
**Supplementary Figure 4.** Full cross-validation and test-set performance of OptiPrime on all datasets examined.
**Supplementary Figure 5.** Full editing outcomes for OptiPrime-assisted correction of *CFTR* mutations.
**Supplementary Figure 6.** Full assessments of editing outcomes for correction of *Kif1a*$^{lgdg}$
**Supplementary Figure 7.** Representative FACS gating strategy for brain nuclei

### Supplementary Text 1. OptiPrime design considerations

#### 1.1. Incorporating mechanistic knowledge into machine learning models

A reaction mechanism describes the process by which a set of reactants are chemically transformed into a set of products. While scientists and engineers can gain a great deal of insight from reaction mechanisms, these insights are typically not readily translatable into high-performance machine learning (ML) models. This deficiency arises because ML models combine information from all of their inputs to generate predictions, learning the relationships between these inputs to approximate the underlying dynamics. While feature engineering can assist with solving this issue, the amount of input data required still grows exponentially relative to the number of features since the model must learn "from scratch" the interactions between features that are relevant and irrelevant.

In developing OptiPrime, we sought to leverage the wealth of knowledge we already had about the PE mechanism. We envisioned representing each possible state that the genomic locus being edited takes during the course of the prime editing (PE) process. Given that there are at most 1–2 PE target loci in each cell, we made the simplifying assumption that the reaction dynamics in PE are governed by pseudo-first order kinetics. Thus, we can write the rate of reaction $i$ at time $t$, which converts genomic state $r$ to genomic state $p$ in the PE mechanism occurring as follows:

$$\mathrm{Rate}_i(t) = -\frac{\mathrm{d}}{\mathrm{d}t}c_r = \frac{\mathrm{d}}{\mathrm{d}t}c_p = k_i c_r(t)$$

where $c_r$ and $c_p$ denote the concentrations of the reactant and product, respectively. However, there can be multiple reactions that result in a given product. If we sum up the rates of all of the reactions that result in state $p$, we find that at time $t$ we have:

$$\frac{\mathrm{d}}{\mathrm{d}t}c_p(t) = \sum_{\substack{j \,\in\, \text{reactions} \\ \text{with product } p}} k_j c_{rj}(t)$$

Writing out the relevant equation for all possible states gives us a system of differential equations that can be rearranged in the form:

$$\frac{\mathrm{d}\mathbf{c}}{\mathrm{d}t} = \mathbf{G}\cdot\mathbf{c}$$

Where $\mathbf{c}(t)$ is a vector representing the concentrations of all states and $\mathbf{G}$ is a "generator" matrix with terms

$$\mathbf{G}_{pr} = \begin{cases} k_{rp} & \text{if } \mathrm{X}_r \to \mathrm{X}_p \text{ is a reaction} \\ -\sum_{q\neq r} k_{rq} & \text{if } r = p \\ 0 & \text{elsewise} \end{cases}$$

where the non-diagonal terms in $\mathbf{G}$ describe the rates at which each reactant turns into a product and the diagonal terms maintain mass balance. We call $\mathbf{G}$ the generator matrix because it takes the same form as the generator matrix for a continuous-time Markov chain.

We can solve this system of differential equations analytically:

$$\mathbf{c}(t) = \exp(\mathbf{G}t) \cdot \mathbf{c}(0); \quad \exp(\mathbf{M}) := \sum_{k=1}^{\infty} \frac{1}{k!}\mathbf{M}^k$$

We noted that the partial derivative of $\exp(\mathbf{G}t)$ with respect to each term in $\mathbf{G}$ is known, and therefore we can train machine learning models that predict individual terms in $\mathbf{G}$ with stochastic gradient descent with backpropagation.

#### 1.2. Mechanistic models used in this work

We modeled the steps in the PE mechanism (as labeled in Fig. S6) as follows:

- $k_{on,PE}$ represents the binding of the PE complex to the target site and subsequent hybridization of the PBS to the nicked flap. $k_{on,PE}$ was modeled with a 4-layer fully connected neural network with 32 hidden neurons in each layer with the following inputs:
  - RNA/RNA duplex melting temperature for the PBS
  - RNA/DNA hybrid melting temperature for the PBS
  - PBS length
  - PBS GC-content
  - The "nick-proximal dinucleotide": 1-hot encoding of spacer positions 16, 17
  - "Total pegRNA concentration": A number between 0 and 1 representing the pegRNA concentration as scaled by the number of poly(T) terminators in the full sequence
  - "Poison pegRNA concentration": A number between 0 and 1 representing the concentration of truncated pegRNAs as scaled by the number of poly(T) terminators in the RTT/PBS
  - "Scaffold defect": $Z - Z_{\text{constrained}}$ where $Z$ is the pegRNA partition function as computed by McCaskill's algorithm and $Z_{\text{constrained}}$ is the partition function when each of the four hairpins in the pegRNA scaffold is constrained. pegRNAs that fold better should have lower scaffold defects.
  - The first (i.e., most scaffold-proximal) base of the RTT
  - Whether the PEmax mutations in Cas9 are present
  - Whether the pegRNA is an epegRNA
  - Whether the pegRNA spacer is 21 bases long
  - Whether the first base of the pegRNA spacer is mismatched
  - The Doench Rule Set 3 score for the pegRNA spacer

  The output of this neural network, $y$, was scaled by the HT-PAMDA score for the PAM sequence with the Cas9 PAM variant used:

  $$\log k_{on,PE} = y + w \cdot \mathbf{P}_{px} + \mathbf{b}_p$$

  where $w$ is a trainable scalar, $\mathbf{P}$ is a constant matrix where term $px$ indexes the HT-PAMDA score of the Cas9 variant $p$ with PAM $x$, and $\mathbf{b}$ is a trainable vector where term $p$ indexes the Cas9 variant.

- $k_{off,PE}$ represents the reverse process of $k_{on,PE}$ and was modeled as a learnable constant.
- $k_{syn}$ represents the **average** rate of addition of each base to the growing $3'$ flap. While we could have modeled $k_{syn}$ as a geometric mean rather than an arithmetic mean, computing the geometric mean is numerically unstable in practice so we chose to use the arithmetic mean instead. We modeled $k_{syn}$ as a linear combination of the number of each base in the RTT along with the RTT secondary structure folding partition function.
- $k_{anneal}$ represents the $3'$ flap annealing into the genome and the excision of the resultant $5'$ flap. Despite attempts to model this step, we found that keeping this rate as a constant was sufficient to achieve the best predictive performance. We speculate that this step was not rate-limiting for any edits in our training set, and that the RTT homology length trends captured by other models are due to differences in DNA repair, rather than annealing itself.
- $k_{on,MutS}$ represents the binding of MutS$\alpha$ or MutS$\beta$ to the heteroduplex. We modeled this rate as a constant since we assume that it is limited by diffusion of the MutS$\alpha$/MutS$\beta$ complexes to the heteroduplex.
- $k_{MMR}$ represents mismatch repair resolving the heteroduplex to the **unedited** sequence. We modeled $k_{MMR}$ as a linear regression with the following inputs:
  - Whether the experiment used MLH1dn
  - The HT-PAMDA score of the PAM sequence for the edited sequence. We included this feature because we hypothesized that re-nicking of the edited strand can occur if a PAM edit is not included.
  - The "nick-proximal dinucleotide" used in $k_{PE,on}$
  - The Doench Rule Set 3 score for the pegRNA spacer
  - The length of homologous sequence downstream of the heteroduplex
  - The "homology-proximal dinucleotide," analogous to the "nick-proximal dinucleotide" but for the last base of the homology arm and the first base downstream of the homology arm.
- $k_{off,MutS}$ represents the unbinding of MutS$\alpha$/MutS$\beta$ to the heteroduplex. This rate is one of the outputs of the HetFormer, described in Note 1.3.
- $k_{rep,u}$ represents the MMR-independent repair of the heteroduplex to the unedited sequence. This rate is one of the outputs of the HetFormer, described in Note 1.3.
- $k_{rep,e}$ represents the MMR-independent repair of the heteroduplex to the edited sequence. This rate is one of the outputs of the HetFormer, described in Note 1.3.

#### 1.3. The HetFormer architecture and pretraining

We sought to create a model that could represent any generic DNA heteroduplex. Although we initially considered using a sequence alignment between the unedited and edited sequences, we found that this representation introduces ambiguities. For example, for the *CFTR* p.F508del mutation, two equally scoring alignments are possible:

```
 I   I   F   G   V
ATC ATC TTT GGT GGT  (ref)
ATC AT- --T GGT GGT  (F508del)
ATC A-- -TT GGT GGT  (F508del)
```

Therefore, the same RTT insertion could encode either a CTT insertion or a TCT insertion. To avoid such ambiguities, we opted to use the HetFormer, which learns the interactions between bases in each strand of the heteroduplex. The HetFormer includes an inductive bias on its cross-attention weights from base pairing probability matrices generated with ViennaRNA.

To pretrain the HetFormer, we algorithmically generated heteroduplexes at random following approximately the ClinVar distribution of pathogenic mutations. We then used the HetFormer to perform simple tasks that we designed:

- Tokens are masked at random ('cloze' style) and the HetFormer must predict identity of the masked tokens
- The HetFormer must classify the heteroduplex as resulting from a substitution, pure insertion, pure deletion, or mix thereof.
- For pure substitutions only, the HetFormer must identify the SNP pairing most prone to MMR. Also, it is regressed on the number of mismatches in the substitution.
- For pure insertions/deletions, the HetFormer must identify the length of insertion/deletion.
- For mixed edit types, the HetFormer is trained to regress on $\log(L_u/L_e)$, which represents the direction and degree to which the length ratio between strands is distorted.

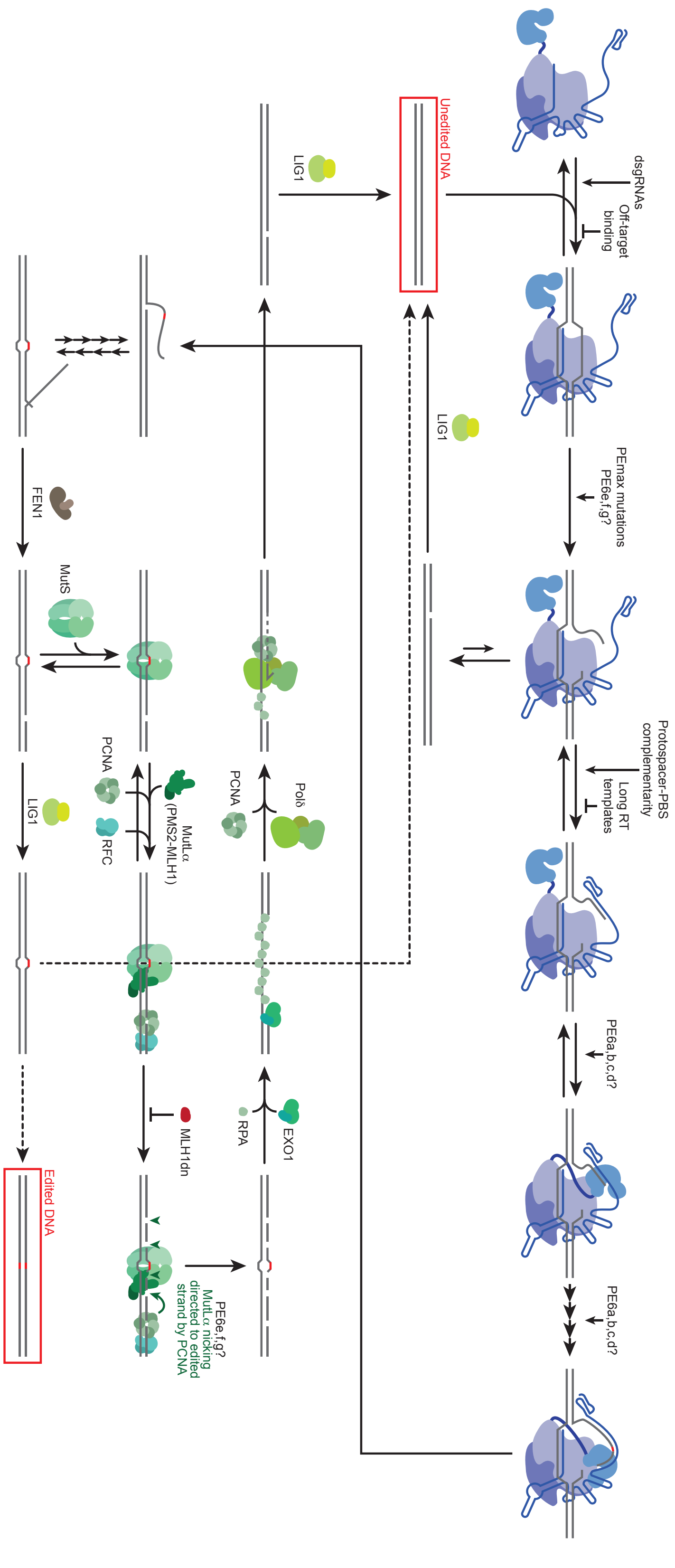


**Supplementary Figure 1**. A mechanism of prime editing in mammalian cells. Standard arrows denote perturbations that speed up a given step, while flathead lines denote those that inhibit a given step.

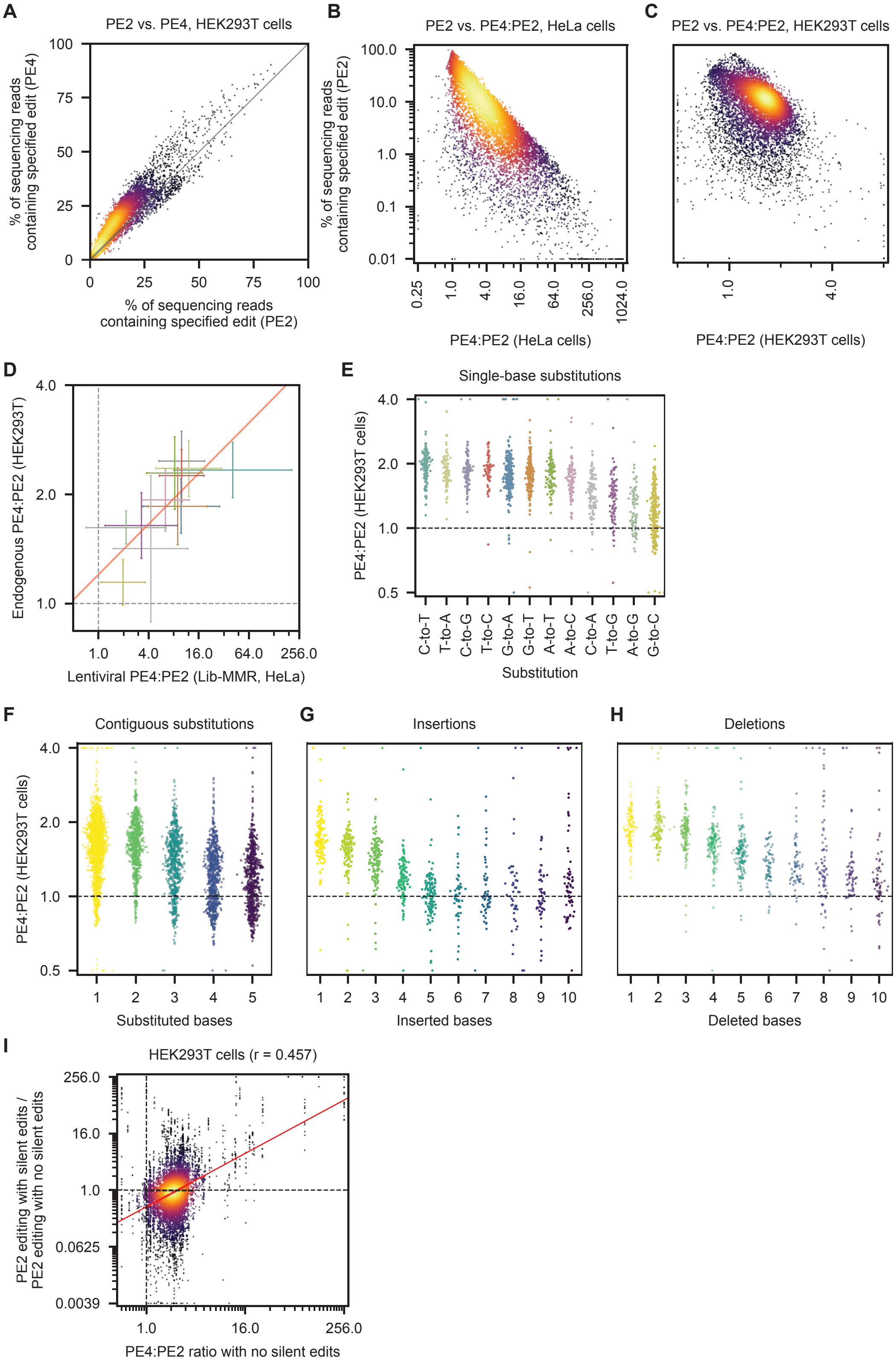


**Supplementary Figure 2.** High-throughput lentiviral screens recapitulate the sequence determinants of mammalian mismatch repair (MMR). **(A)** A scatter plot comparing prime editing efficiencies across Lib-MMR between PE2 (*x*-axis) and PE4 (*y*-axis). The gray line plots the equation *y* = *x*. PE4 somewhat outperforms PE2 in HEK293T cells. **(B–C)** Scatter plots comparing PE4:PE2 ratios (*x*-axis, log-scale) and PE2 editing efficiency (*y*-axis, log-scale) in (B) HeLa cells and (C) HEK293T cells.

PE4:PE2 correlates better with PE2 in HeLa cells than in HEK293T cells. **(D)** A scatter plot comparing PE4:PE2 ratios for single-base substitutions in the lentiviral Lib-MMR context against those previously obtained at endogenous loci[1]. Colors correspond to those in Figures 2E and S2E. Error bars represent ± 1 standard deviation. The red line plots the total least squares regression line of the mean PE4:PE2 ratios. **(E)** Beeswarm plots of PE4:PE2 in HEK293T cells for all twelve possible single-base substitutions, sorted by median PE4:PE2. See also Figure 2E. **(F–H)** Beeswarm plots comparing PE4:PE2 in HEK293T cells across varying lengths of (F) contiguous substitutions, (G) pure insertions, and (H) pure deletions. See also Figure 2F–H. **(I)** A scatter plot comparing PE4:PE2 editing efficiency ratio of each edit to the ratio of PE2 editing efficiency with and without silent edits across Lib-CV in HEK293T cells. Improvements in PE2 efficiency from including silent edits are positively correlated with PE4:PE2 efficiency ratios of the reference edit without any silent edits in HeLa cells. The red line is a total least squares regression line. Brighter colors correspond to increased point density.

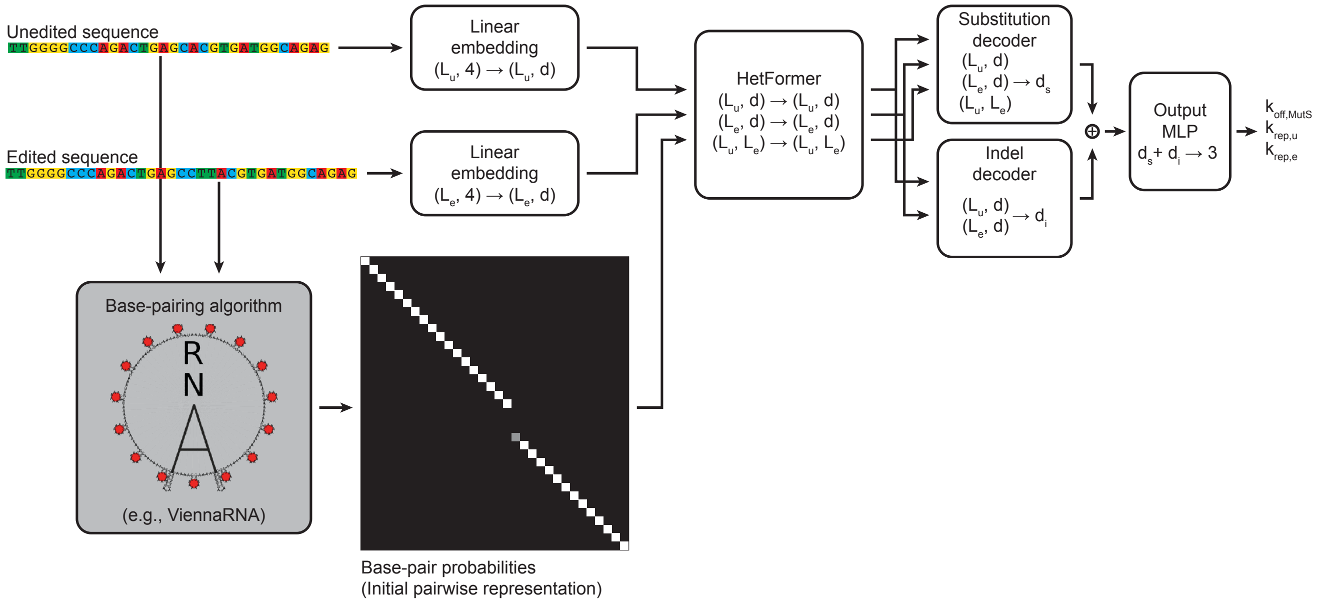


**Supplementary Figure 3**. The HetFormer is a neural network designed to model DNA heteroduplexes. The HetFormer is a deep neural network architecture that combines sequence information with base-pairing probability information about a heteroduplex to predict pseudo-rates relevant for DNA repair.

**A**

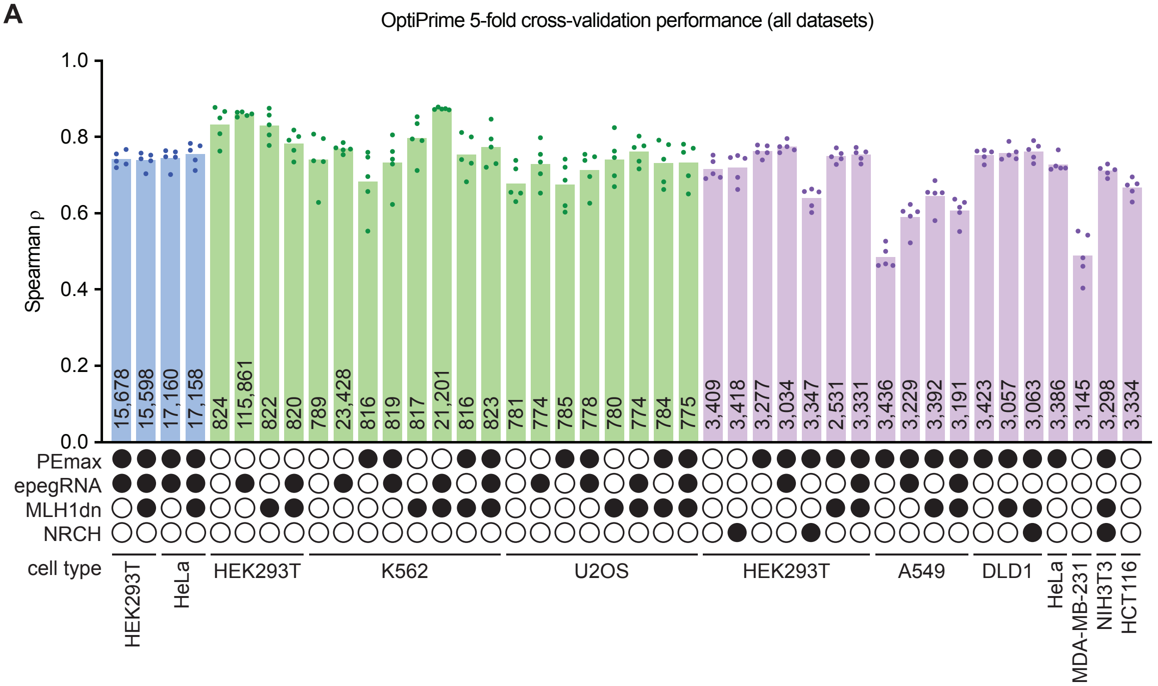
OptiPrime 5-fold cross-validation performance (all datasets)
Spearman ρ
PEmax
epegRNA
MLH1dn
NRCH
cell type
HEK293T
HeLa
HEK293T
K562
U2OS
HEK293T
A549
DLD1
HeLa
MDA-MB-231
NIH3T3
HCT116
15,678
15,598
17,160
17,158
824
115,861
822
820
789
23,428
816
819
817
21,201
816
823
781
774
785
778
780
774
784
775
3,409
3,418
3,277
3,034
3,347
2,531
3,331
3,436
3,229
3,392
3,191
3,423
3,057
3,063
3,386
3,145
3,298
3,334

**B**

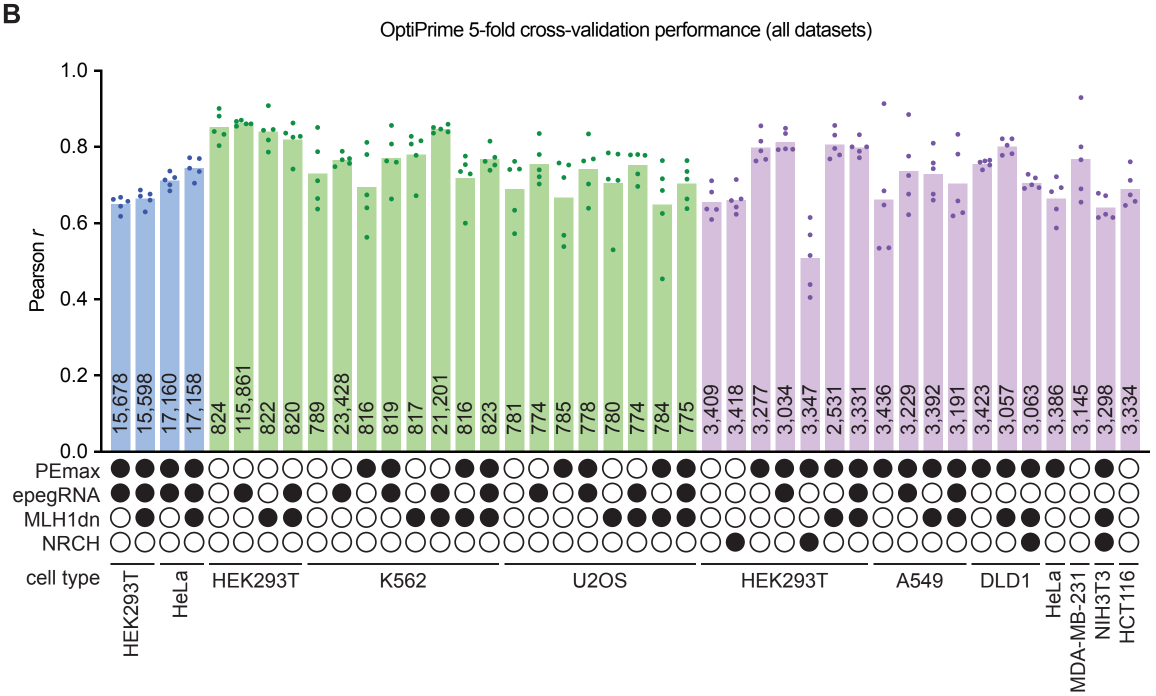
OptiPrime 5-fold cross-validation performance (all datasets)
Pearson r
PEmax
epegRNA
MLH1dn
NRCH
cell type
HEK293T
HeLa
HEK293T
K562
U2OS
HEK293T
A549
DLD1
HeLa
MDA-MB-231
NIH3T3
HCT116

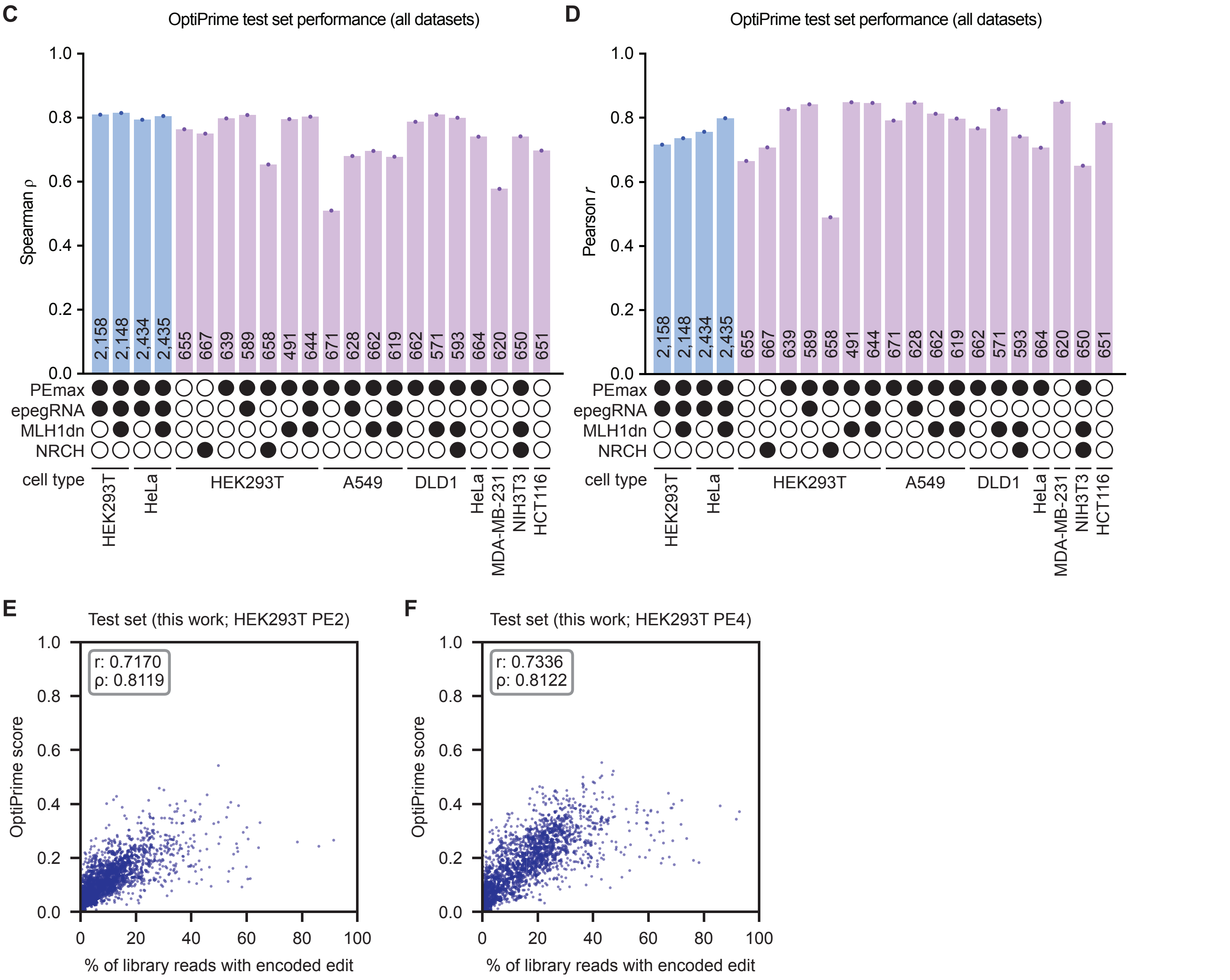


**Supplementary Figure 4.** Full cross-validation and test-set performance of OptiPrime on all datasets examined. **(A, B)** Bar graphs showing OptiPrime performance on training sets under 5-fold cross-validation evaluated with either (A) Spearman ρ or (B) Pearson *r*. Filled circles in *x*-axis labels depict the specific PE system used for each dataset (PE2 vs. PEmax protein, epegRNA vs. pegRNA, PE2 vs PE4 system, SpCas9 or SpCas9-NRCH). Blue bars are datasets collected in this work, green bars are datasets collected in Mathis et al.[2, 3], and purple bars are datasets collected in Yu et al.[4]. Numbers overlaid on each bar represent the total number of data points in each training set. **(C, D)** Bar graphs showing OptiPrime performance on lentiviral library-derived test sets evaluated with either (A) Spearman ρ or (B) Pearson *r*. Note that the datasets from Mathis et al.[2, 3] were evaluated with nested cross-validation rather than held-out test sets, and thus are not included in this comparison. (C) Spearman ρ or (B) Pearson *r* values for each training dataset. Filled circles in *x*-axis labels depict the specific PE system used for each dataset (PE2 vs. PEmax protein, epegRNA vs. pegRNA, PE2 vs PE4 system, SpCas9 or SpCas9-NRCH). Blue bars are datasets collected in this work and purple bars are datasets collected in Yu et al.[4]. Numbers overlaid on each bar represent the total number of data points in each test set. **(E, F)** Scatter plot of test set predictions with OptiPrime for the data collected in this study in HEK293T cells with (E) PE2 and (F) PE4.

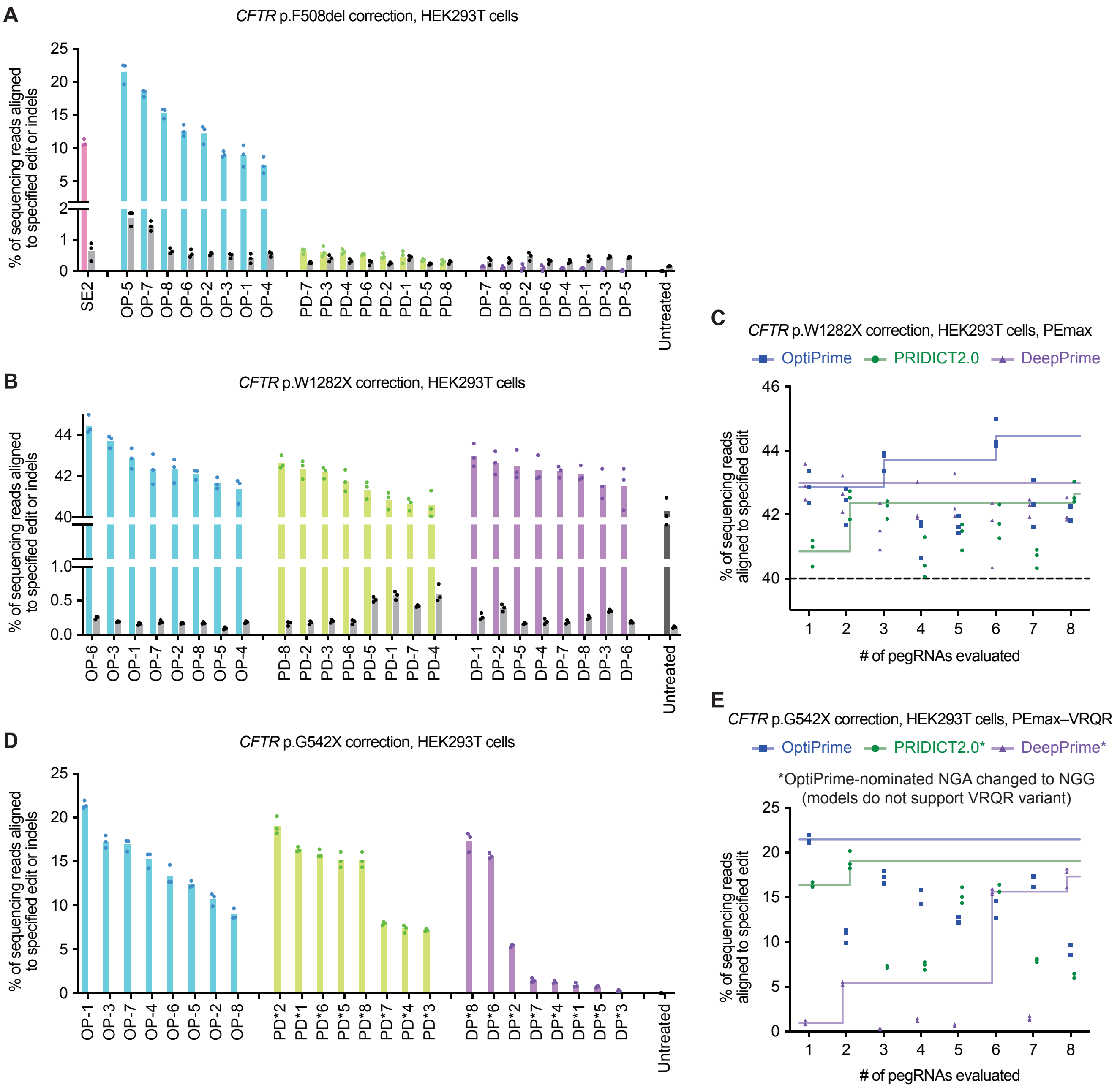


**Supplementary Figure 5.** Full editing outcomes for OptiPrime-assisted correction of *CFTR* mutations. **(A)** Full editing data for the pegRNAs displayed in Figure 6C for the correction of *CFTR* p.F508del. **(B)** Correction of the *CFTR* p.W1282X mutation with OptiPrime, PRIDICT2.0, and DeepPrime in HEK293T cells. Points denote individual experimental editing efficiencies (*N* = 3) for each ranked pegRNA on the *x*-axis, while lines indicate the maximum editing efficiency achieved after evaluating the top *x*-ranked pegRNAs. **(C)** Full editing data for the pegRNAs displayed in Figure S6B for the correction of *CFTR* p.W1282X. **(D)** Correction of the *CFTR* p.G542X mutation with OptiPrime, PRIDICT2.0, and DeepPrime in HEK293T cells. PRIDICT2.0 and DeepPrime do not offer support for the Cas9-VRQR PAM variant, so the input allele was modified from NGA to NGG for PAM compatibility and the corresponding RTT/PBS length outputs were used. Points denote individual experimental editing efficiencies (*N* = 3) for each ranked pegRNA on the *x*-axis, while lines indicate the maximum editing efficiency achieved after evaluating the top *x*-ranked pegRNAs. **(E)** Full editing data for the pegRNAs displayed in Figure S6D for the correction of *CFTR* p.W1282X.

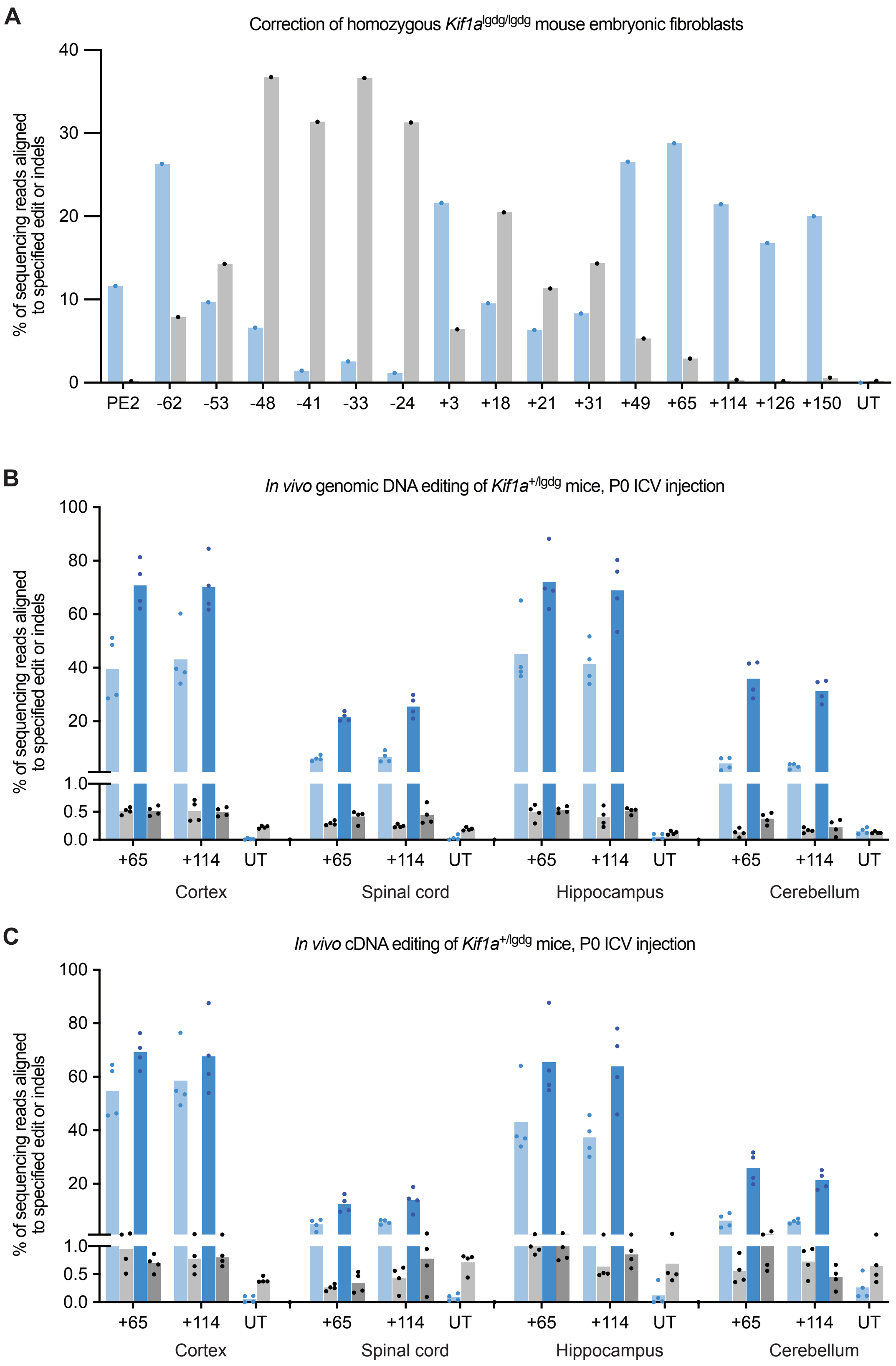


**Supplementary Figure 6.** Full assessments of editing outcomes for correction of *Kif1a*$^{lgdg}$. **(A)** Prime editing efficiency and indel formation rates for nicking guide strategies in homozygous *Kif1a*$^{lgdg/lgdg}$ mouse embryonic fibroblasts. **(B, C)** *In vivo* prime editing efficiency in (B) genomic DNA or (C) cDNA were evaluated from both bulk tissue and from GFP-positive nuclei (enriched for AAV-transduced cells).

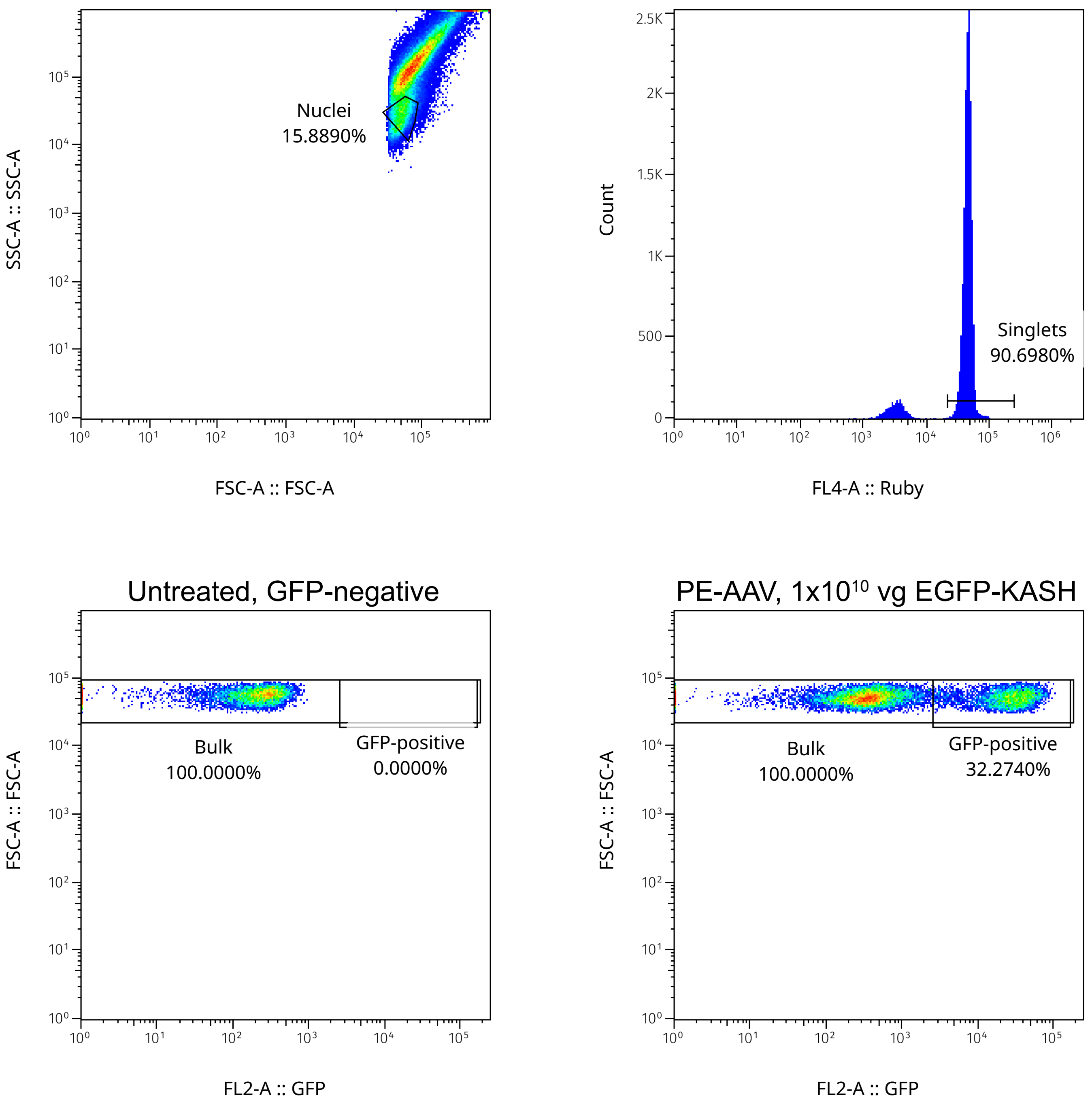


**Supplementary Figure 7.** Representative FACS gating strategy for brain nuclei. Singlet DyeCycle Ruby-positive events were collected and back-gated to establish forward and side scatter area gates for nuclei. Nuclei were then sorted based on GFP fluorescence to yield “bulk” (containing all nuclei irrespective of GFP expression) and “GFP-positive” populations.
